# Functional divergence shaped the network architecture of plant immune receptors

**DOI:** 10.1101/2023.12.12.571219

**Authors:** Ching-Yi Huang, Yu-Seng Huang, Yu Sugihara, Hung-Yu Wang, Lo-Ting Huang, Juan Carlos Lopez-Agudelo, Yi-Feng Chen, Kuan-Yu Lin, Bing-Jen Chiang, AmirAli Toghani, Jiorgos Kourelis, Lida Derevnina, Chih-Hang Wu

**Affiliations:** Institute of Plant and Microbial Biology, Academia Sinica, Taipei, Taiwan; The Sainsbury Laboratory, University of East Anglia, Norwich Research Park, Norwich, UK; Crop Science Center, Department of Plant Sciences, University of Cambridge, Cambridge, UK

**Keywords:** plant immunity, helper NLR, NRC network, sensor-helper compatibility, subfunctionalization, transient interactions, TurboID

## Abstract

In solanaceous plants, several sensor NLRs and their helper NLRs, known as NRC, form a complex network to confer immunity against pathogens. While the sensor NLRs and downstream NRC helpers display diverse genetic compatibility, the evolution and molecular basis of the complex network structure remained elusive. Here we demonstrated that functional divergence of NRC3 variants has shaped the genetic architecture of the NLR network. Natural NRC3 variants form three allelic groups displaying distinct compatibilities with sensor NLRs. Ancestral sequence reconstruction and analyses of natural and chimeric variants identified six key amino acids involved in sensor-helper compatibility, with two residues critical for subfunctionalization. Co-functioning Rpi-blb2 and NRC3 variants showed stronger transient interactions upon effector detection, with NRC3 membrane-associated complexes forming subsequently. Our findings reveal how mutations in helper NLRs, particularly NRC3, have driven the evolution of their transient interactions with sensor NLRs, leading to subfunctionalization and contributing significantly to the complexity of the NRC network in plant immunity.

**Teaser:** Helper NLR subfunctionalization alters transient interactions with sensor NLRs, enhancing plant immune system complexity.

## Introduction

NLR (nucleotide-binding domain and leucine-rich repeat) proteins are intracellular receptors used by both plants and animals to detect invading pathogens (*1*, *2*). They usually consist of an N-terminal domain that is essential for initiating downstream responses, an NB-ARC domain that hydrolyzes ATP, and a leucine-rich repeat region that is involved in pathogen recognition (*2*). Typical plant singleton NLRs, such as ZAR1, can recognize pathogen molecules and initiate downstream responses through the formation of the resistosome complex that functions as a calcium channel (*3–5*). However, some NLR proteins have evolved into helper and sensor NLRs that function together, forming complex NLR networks to confer resistance against invading microbes (*2*, *6*, *7*). In mammals, NLRC4 is a typical helper NLR that forms inflammasome complexes with the sensor NLRs NAIP2 or NAIP5 (*8*, *9*). In plants, the NRC (NLR-required for cell death) family and the ADR1/NRG1 family represent two groups of helper NLRs functioning downstream of different sets of sensor NLRs (*10–12*).

The NRC network of solanaceous plants is composed of multiple sensor NLRs that detect molecules from various pathogens and a few NRC helper NLRs that are essential for mediating cell death responses for the sensor NLRs (*10*). Three of the NRCs, namely NRC2, NRC3, and NRC4, display partial genetic redundancy as well as specificity to different sensor NLRs, resulting in an NLR network with intricate genetic structure. For example, the sensor NLR Rpi-blb2 signals through NRC4 but not NRC2 and NRC3 in the model solanaceous plant species *Nicotiana benthamiana*, the sensor NLR Prf signals through NRC2 and NRC3, whereas the sensor NLR Rx can signal redundantly through NRC2, NRC3 or NRC4 (*10*). Recent studies indicate that NRC0 is the only conserved NRC across asterids plants (*13*). NRC0 orthologs are often located in a gene cluster together with the sensor NLRs that are NRC0-dependent (*13*, *14*). In different lineages of asterids, members of the NRC0 subclades are partially interchangeable, indicating that NRC0 orthologs have functionally diversified in different species (*13*, *14*). The NRC networks likely originated from a sensor-helper NLR pair that emerged predating the divergence of asterids and Caryophyllales, and massively expanded in lamiids, in particular in solanaceous plants and several *Ipomoea* species (*10*, *13*, *14*).

While NLRs often show high diversity across plant species, many well-studied examples consist of NLRs that have remained largely conserved across evolution. For example, ZAR1, which represents an ancient category of plant immune receptors, indirectly recognizes effectors by engaging with its RLCK (Receptor-Like Cytoplasmic Kinase) partners, forming pentameric resistosome complexes associated with the membrane (*3*, *15*). These complexes serve as calcium-permeable channels to trigger immune responses (*5*). Similarly, helper NLRs, including ADR1, NRG1, and NRCs form membrane-associated punctate and high molecular weight complexes upon activation by their respective sensor NLRs (*16–20*). While the NRC family members are broadly conserved functioning as helper NLRs, the NRC networks have diversified across different asterids species (*13*, *14*). As the sensor and helper NLRs within the NRC network trace back to a common ancestral helper-sensor cluster similar to other NLR pairs, it is intriguing to observe that diverse levels of compatibility have evolved among multiple helper NLRs and sensor NLRs (*13*, *14*). However, how the helper-sensor NLR compatibility is determined and evolved in the network is not clear.

In this study, we addressed a fundamental feature of the NRC network architecture: what are the molecular determinants of sensor/helper specificity? We focused on the evolution of specificity between NRC3 orthologs to the sensor NLR Rpi-blb2. We found that Rpi-blb2 can only signal (referred to as compatible from here on) through a subset of NRC3 variants. Using ancestral sequence reconstruction, we showed that the change of compatibility evolved through subfunctionalization. We mapped the determinants that affect the compatibility of NRC3 orthologs with Rpi-blb2 to three residues in the NB-ARC domain and three residues in the LRR domain. These residues contribute to a transient interaction between the sensor and helper NLR proteins upon effector detection. Furthermore, the transient interaction was detected before the formation of the activated membrane-associated complexes of NRC3. We propose that the NRC network evolved through successive cycles of helper NLR duplications followed by mutations leading to subfunctionalization. These subfunctionalization events may divide the NLR network into smaller subnetworks, increasing the complexity of the immune system.

## Results

### NRC3 orthologs and paralogs show different compatibility to the sensor NLR Rpi-blb2

To gain insights into the molecular mechanisms and evolution of the NRC network, we cloned several NRC homologs from tomato and *N. benthamiana* and performed complementation assays with several sensor NLRs in the *nrc2/3/4* CRISPR knockout (*nrc2/3/4*_KO) *N. benthamiana* line (*21*). We reasoned that exploring closely related NRC homologs that show different compatibilities to the sensor NLRs may shed light on the molecular determination and evolution of the helper-sensor compatibility. The cloned NRCs were grouped into NRC0 to NRCX based on the phylogenetic analysis (**Fig. S1A**) (*10*, *22–24*). We individually expressed these NRCs with Rx, Sw5b, Prf (Pto/AvrPto), Gpa2, Rpi-blb2, and R1 with their corresponding effector proteins in *N. benthamiana* leaves, and performed cell death intensity quantification using autofluorescence-based imaging **(Fig. S2A**). While most of the sensor-helper genetic dependency results were consistent with the previous report, tomato NRC3 (SlNRC3) but not *N. benthamiana* NRC3 (NbNRC3) rescued Rpi-blb2-mediated cell death in the *nrc2/3/4*_KO *N. benthamiana* (**Fig. 1A-B and Fig. S2B-F**) (*10*). To explore the differences among the NRC3 variants, we performed phylogenetic analyses of several NRC3 sequences identified from solanaceous plants. We found that the NRC3 homologs are clustered into three allelic groups, including Group A (NRC3a) which contains orthologs of NRC3 from *Solanum* and *Capsicum* species, and Group B (NRC3b) and Group C (NRC3c) which contains NRC3 from *Nicotiana* species (**Fig. 1C**). Both *N. tabacum* and *N. sylvestres* harbor sequences of NRC3b and NRC3c, whereas *N. benthamiana* only contains one NRC3c sequence (**Fig. 1C**). We cloned several of these NRC3 variants and tested their ability to rescue Rpi-blb2, Prf (Pto), and Rx-mediated cell death in the *nrc2/3/4*_KO *N. benthamiana.* We found that the two NRC3a variants tested were able to rescue Rpi-blb2, Prf, and Rx-mediated cell death, while the NRC3b variants failed to rescue cell death mediated by these sensors. Interestingly, most NRC3c variants were able to function with Prf and Rx but not Rpi-blb2 **(Fig. 1D**). The accumulations of all the natural variants were detectable, and none of them showed strong auto-activity in inducing cell death when expressed alone (**Fig. 1E and Fig. S3**). These results suggested that functional divergence has evolved among the NRC3 homologs.

**Fig. 1.**
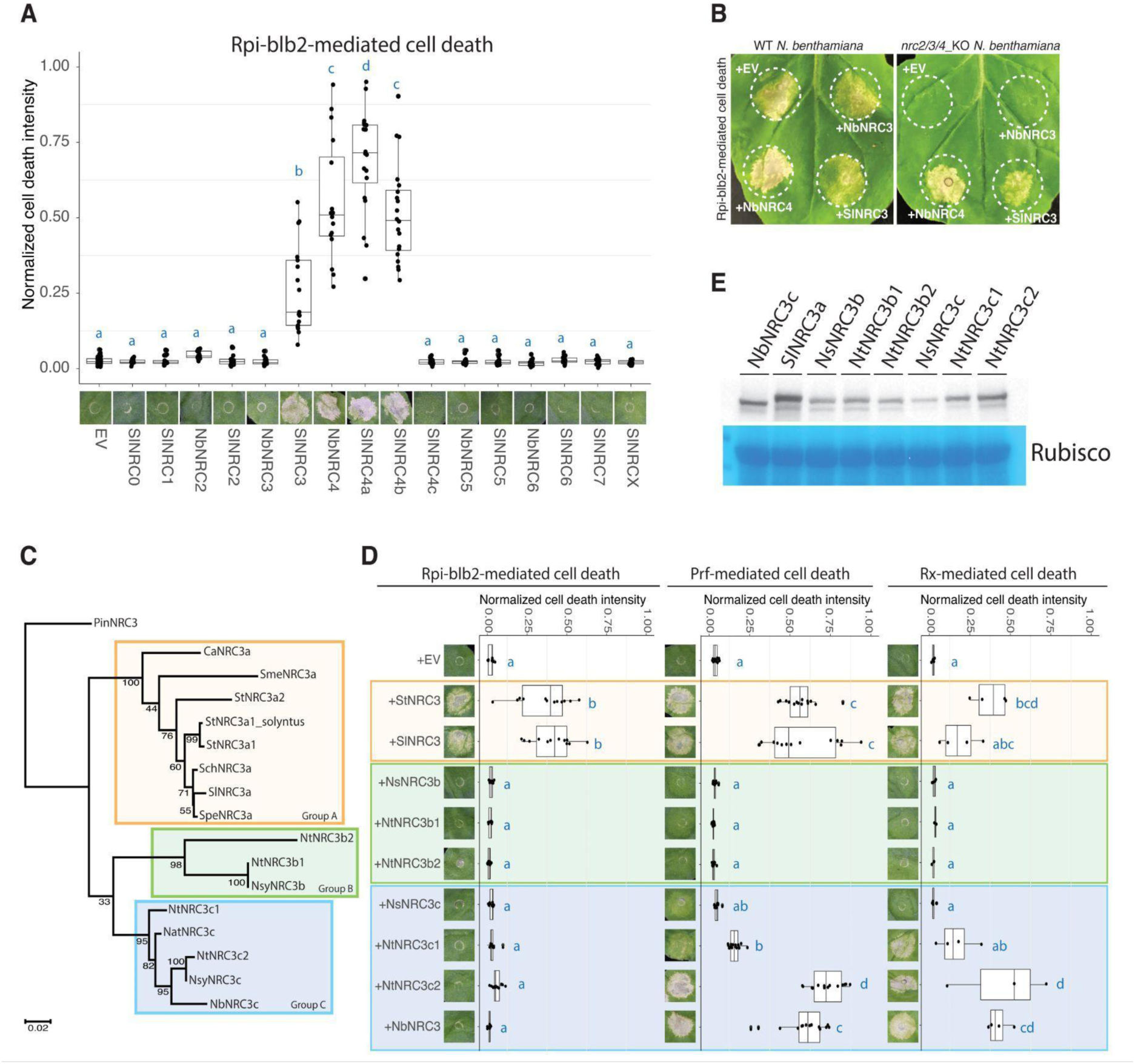
Rpi-blb2 signals through NRC3a but not NRC3b or NRC3c. (A) Cell death assays of Rpi-blb2 with different NRCs. Rpi-blb2 and AVRblb2 were co-expressed with indicated NRCs cloned from tomato and *N. benthamiana* in *nrc2/3/4*_KO *N. benthamiana.* Cell death intensity and phenotypes were recorded at 6 dpi. (B) Assays of Rpi-blb2-mediated cell death rescued by NRC variants. Rpi-blb2 and AVRblb2 were co-expressed with NRCs as indicated in both WT or *nrc2/3/4*_KO *N. benthamiana*. (C) Phylogenetic analysis of NRC3 natural variants identified from tomato, tobacco potato, pepper, and eggplant. Sequence alignment of the NB-ARC domain was used to generate the phylogenetic tree using the Maximum likelihood method with 100 bootstrap tests. Petunia NRC3 (PinNRC3) was selected as an outgroup. The orange, green, and blue boxes indicate allelic groups A, B, and C, respectively. (D) Cell death assays of different sensor NLRs with NRC3 natural variants. The cloned NRC3 natural variants were co-expressed with Rpi-blb2/AVRblb2, Pto/AvrPto, or Rx/CP in *nrc2/3/4*_KO *N. benthamiana*. (E) Protein accumulation of NRC3 natural variants. NRC3 natural variants were transiently expressed in WT *N. benthamiana.* The proteins were extracted from leaf samples at 2 dpi and the NRC3 protein accumulations were detected by α-myc antibody. SimplyBlue SafeStain-staining of Rubisco was used as the loading control. The dot plots in (A) and (D) represent cell death intensity quantified by UVP ChemStudio PLUS at 6 dpi. Statistical differences were analyzed with Tukey’s HSD test (p<0.05).

### Ancestral reconstructions reveal subfunctionalization of NRC3c towards loss of compatibility with Rpi-blb2

To determine what functional divergence events occurred during the evolution of NRC3, we performed functional assays of reconstructed ancestral NRC3 variants. We extracted 324 non-redundant nucleotide sequences of the NRCX, NRC1, NRC2, and NRC3 clades from 124 solanaceous genomes, and then used FastML to reconstruct the ancestral NRC sequences (**Fig. S4A**). We synthesized the full-length of five ancestral NRC3 variants reconstructed from the FastML, including N4, N95, N89, N88, and N3 that represent the ancestral state of NRC3a, NRC3b, NRC3c, NRC3b/c, and before the divergence of the three allelic groups (**Fig. 2A, Fig. S5**). We then tested the degree to which these ancestral variants can rescue Rpi-blb2, Prf (Pto), and Rx-mediated cell death. We found that while N3, N4, and N88 rescue Rpi-blb2-mediated cell death, the ancestral variants N95 (NRC3b) and N89 (NRC3c) show no or little activity in rescuing Rpi-blb2-mediated cell death (**Fig. 2B**). While most of these ancestral variants (N3, N4, N88, and N89) rescued Prf (Pto) and Rx-mediated cell death, N95 was the only variant that failed to rescue any cell death tested (**Fig. 2B**). The accumulations of all of these ancestral variants were detectable with low or no auto-activity (**Fig.e S6**). We introduced a D to V mutation into the MHD motif of N95 and NRC3b variants, and found that none of these variants induce cell death in *N. benthamiana*, suggesting that this group of NRC3 has nonfunctionalized during the evolutionary process (**Fig. 2C**). These results indicate that NRC3 in the ancestral species likely functions together with Rpi-blb2/Prf/Rx, whereas the NRC3 variants that evolved in *Nicotiana* species acquired mutations leading to nonfunctionalization (NRC3b) or subfunctionalization (NRC3c), losing their ability to work together with Rpi-blb2.

**Fig. 2.**
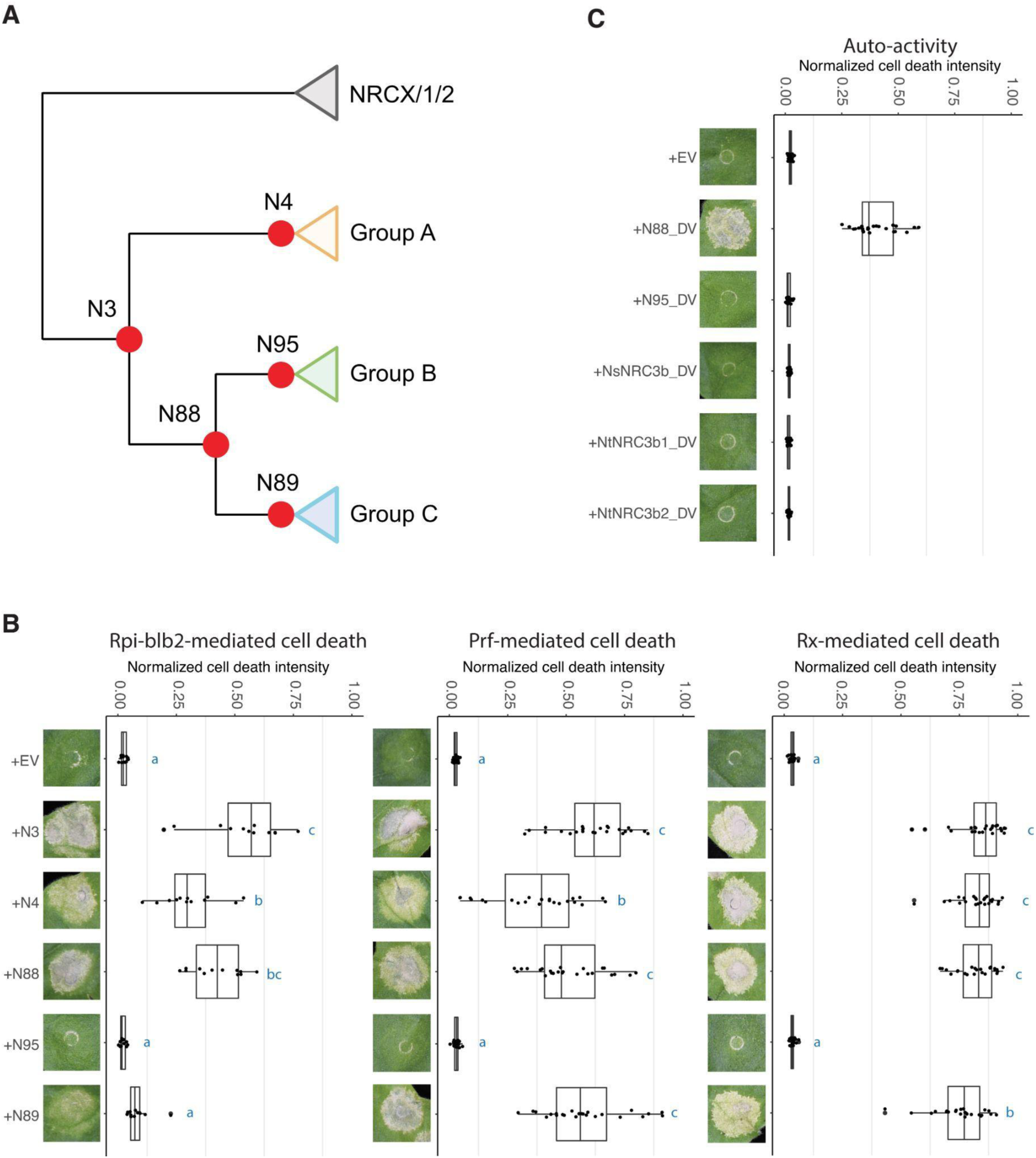
Subfunctionalization contributes to the evolution of NRC3c. (A) Phylogenetic tree of NRC3 natural variants. Orange, green, and blue boxes represent allelic groups A, B, and C, respectively. Red dots indicate the nodes of reconstructed ancestral NRC3 variants. (B) Cell death assays of ancestral NRC3 variants. The ancestral variants were co-expressed with Rpi-blb2/AVRblb2, Pto/AvrPto, or Rx/CP in *nrc2/3/4*_KO *N. benthamiana*. (C) Cell death analysis of NRC3b_DV, N88_DV and N95_DV. The NRC3_DV variants carry a D to V mutation in the MHD motif. These variants were expressed alone in WT *N. benthamiana*. The dot plots represent cell death intensity quantified by UVP ChemStudio PLUS at 6 dpi. Statistical differences were analyzed with Tukey’s HSD test (p<0.05).

To further test this hypothesis, we synthesized NRC3 of *Petunia inflata* (PinNRC3). The PinNRC3 from the genome database contains an indel of 9 amino acids between the CC and NB-ARC domains. Therefore, we manually curated the indel by using the sequences from SlNRC3a, and found that this manually curated PinNRC3 variant is able to rescue all three cell death phenotypes tested (**Fig. S7**). Taken together, these results support the hypothesis that the ancestral NRC3 can function with a broader collection of sensor NLRs, while NRC3c variants subfunctionalized to work with a smaller subset of sensor NLRs.

### Six NRC3 amino acid residues contribute to the compatibility of NRC3 with Rpi-blb2

Since SlNRC3a and NbNRC3c showed robust differences in their ability to function with Rpi-blb2, we focused on these two NRC3 variants to identify the residues that affect their compatibility with Rpi-blb2. We reasoned that differences in amino acids at these positions may infer the mutations that occurred during NRC3c subfunctionalization. Pairwise sequence comparison indicated that the two NRC variants share around 86.42% overall sequence identity, with 87%, 90%, and 83% for the CC domain, NB-ARC domain, and LRR domain, respectively (**Fig. S8A**). We then generated chimeric proteins between the two NRC3 variants and determined which regions contribute to the compatibility with Rpi-blb2. These constructs were named NNS, NSN, NSS, SSN, SNS, and SNN (N stands for *N. benthamiana*; S stands for *S. lycopersicum*) (**Fig. S8B**). The chimeric variants carrying the CC domain from SlNRC3a (SNS, SSN, SNN) showed no or low activity in rescuing Rpi-blb2-mediated cell death, suggesting that the CC domain is not critical in determining their compatibility with Rpi-blb2 (**Fig. S8B**). While the chimeric variants NSN and NNS showed low activities, the variant NSS fully rescued Rpi-blb2-mediated cell death, similar to the degree observed using SSS (SlNRC3) (**Fig. S8B**). None of these chimeric NRC3 variants were auto-active, with SNS being the only variant that failed to rescue Prf-mediated cell death (**Fig. S8B**). All of these NRC3 chimeric variants were detectable using immunoblot analysis (**Fig. S8C**). These results suggest that the NB-ARC and LRR domains cooperatively determine the compatibility of NRC3 variants with Rpi-blb2.

To further identify the key polymorphisms within the NB-ARC and LRR domains that contribute to helper-sensor compatibility, we generated two sets of chimeric variants of Nb/SlNRC3 for complementation assays, focusing on the polymorphisms in either the NB-ARC or the LRR domain. We divided the NB-ARC domain into 10 smaller regions based on sequence alignment and generated corresponding chimeric NRC3 variants named NN_1-10_S (**Fig. 3A and Fig. S9**). While most of the NN_1-10_S chimeric NRC3 variants showed low activity in rescuing Prf-mediated cell death, NN_3_S was the only variant that fully rescued both Rpi-blb2- and Prf-mediated cell death (**Fig. 3A and Fig. S9**). This region contained six amino acid differences between the two variants (**Fig. 3A**). To identify the regions in the LRR domain that contribute to helper-sensor compatibility, we generated chimeric protein NSN_1-5_, with polymorphisms from the LRR of tomato NRC3 swapped into the NSN background (**Fig. 3B**). While the NRC3 variant NSN_1_ was autoactive (**Fig. S10A**), the variant NSN_2_ and NSN_5_ both partially rescued Rpi-blb2-mediated cell death in the *nrc2/3/4*_KO *N. benthamiana* (**Fig. 3B and Fig. S10**). Thus, we generated another chimeric NRC3 variant, named NSN_25_, and found that this variant fully rescued Rpi-blb2 cell death (**Fig. 3B**). These results suggest that both the NB-ARC and LRR domains cooperatively enable the NRC3 variant to function with Rpi-blb2. To pinpoint the residues in region 2 of the LRR domain, we tested additional chimeric NRC3 variants. We found that a single amino acid change (I to T) at position 642 is sufficient to enable NSN to weakly rescue Rpi-blb2-mediated cell death (**Fig. S11A**). This single amino acid change also conferred full activity in rescuing Rpi-blb2 cell death in the NSN_5_ background (**Fig. 3C and Fig. S11B**). We divided region 5 of the LRR domain into five fragments and generated variants named NSN_5a-e_, and found that NSN_5b_ and NSN_5e_ partially rescue Rpi-blb2-mediated cell death (**Fig. S12A and C**). We combined these polymorphisms into the NSN_2_ background to generate a new variant named NSN_25be_. As expected, the NRC3 variant NSN_25be_ fully rescued Rpi-blb2-mediated cell death (**Fig. 3D and Fig. S12B and D**).

**Fig. 3.**
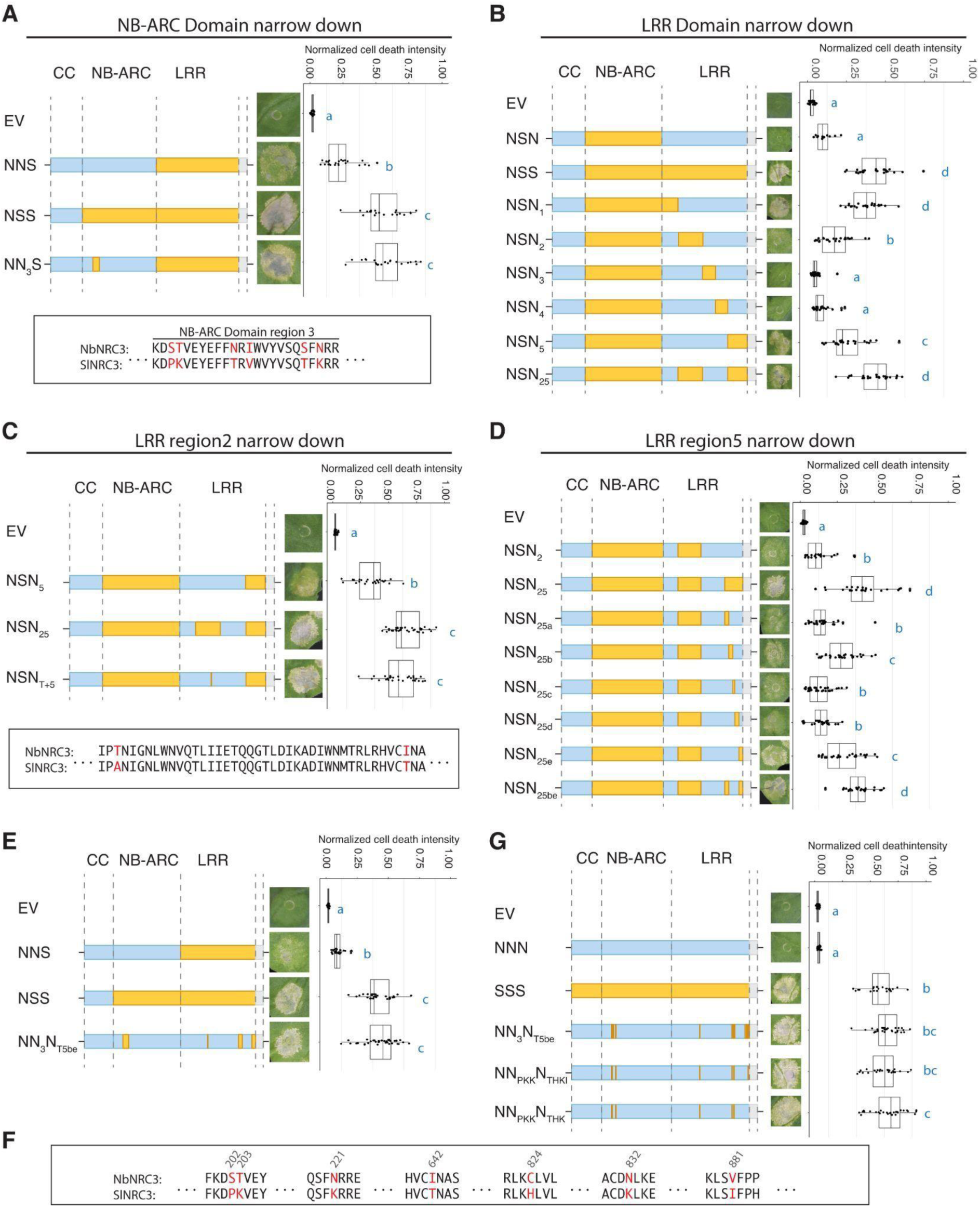
Changing six residues in NbNRC3 enabled it to function with Rpi-blb2. Cell death assays of chimeric NRC3 variants designed to investigate polymorphisms in (A) the NB-ARC domain, (B) the LRR domain, (C) LRR domain region 2, and (D) LRR domain region 5 that contribute to the helper-sensor compatibility. (E) Cell death assays of chimeric NRC3 variants that carry polymorphisms identified in (A) to (D). (F) Polymorphisms of NbNRC3 and SlNRC3 at the seven positions identified (highlighted in red). (G) Cell death assays of chimeric NRC3 variants NN_PKK_N_THKI_ and NN_PKK_N_THK_. Cell death assays were performed by co-expressing chimeric NRC3 variants with Rpi-blb2/AVRblb2 in *nrc2/3/4*_KO *N. benthamiana*. The dot plots represent cell death intensity quantified by UVP ChemStudio PLUS at 6 dpi. Statistical differences were analyzed with Tukey’s HSD test (p<0.05).

We sought to pinpoint the amino acid residues in each region that enabled NRC3 variants to function with Rpi-blb2 in the NbNRC3c background. We introduced the polymorphisms identified above into NbNRC3c, generating a chimeric variant named NN_3_N_T5be_. We found that NN_3_N_T5be_ can fully rescue cell death mediated by Rpi-blb2 in *nrc2/3/4*_KO *N. benthamiana* (**Fig. 3E and Fig. S13**). Through additional sets of loss-of-function screens, we found that amino acid changes of S202P, T203K, N221K, C824H, N832K, and V881I quantitatively affect the ability of NRC3 to function with Rpi-blb2 (**Fig. S14 and Fig. 3F**). Thus, we introduced these mutations into NbNRC3c together with I642T, and found that this new NRC3 variant, named NN_PKK_N_THKI_, can fully rescue Rpi-blb2-mediated cell death (**Fig. 3G and Fig. S15**). Since V and I at position 881 are similar residues, we further generated NN_PKK_N_THK_ and found that this variant showed similar activity to NN_PKK_N_THKI_ (**Fig. 3G**). Based on the above results, we concluded that six amino acid changes of 6 amino acids, three in the NB-ARC domain and three in the LRR domain, enable NbNRC3c to function with Rpi-blb2. These results suggest that changes in amino acids at these positions pose a major contribution to the helper-sensor compatibility and subfunctionalization during the evolution of the NRC3c allelic group.

### Two K to N mutations in the NB-ARC and LRR domains play critical roles in NRC3 subfunctionalization

Next, we looked into the polymorphism of these 6 positions in the NRC3 natural variants tested above (**Fig. 1C**). We found that all the NRC3 variants from allelic group A possess PKKTHK, and all the variants from allelic group B and the outgroup PinNRC3 possess PKKTRK (**Fig. 4A and Fig. S4B**). Interestingly, sequences from allelic group C showed higher diversity, with NbNRC3c being the most diverse variant (**Fig. 4A**). We introduced these different polymorphisms into the NbNRC3c background and tested the ability of these variants to rescue Prf and Rpi-blb2 cell death in *nrc2/3/4*_KO *N. benthamiana*. While all these variants rescued Prf-mediated cell death, NRC3 variants with PKKTHK or PKKTRK, but not PKNTRN or PTNTRN, were fully able to rescue Rpi-blb2-mediated cell death (**Fig. 4A and Fig. S16A**). Consistent with the results from the polymorphisms found in the natural variants, the major differences between N88 (PKKTRK) and N89 (PKNTRN) were also due to two K to N mutations (**Fig. 4B**). These findings indicate that the two mutations converting K to N are likely the most crucial changes during the subfunctionalization process. To further test this hypothesis, we introduce the two K to N mutations into the ancestral variants N88 (N88^NN^) or the N to K mutations into the ancestral variants N89 (N89^KK^). We found that N88^NN^ showed reduced activity and N89^KK^ showed increased activity compared to their ancestral states respectively (**Fig. 4B and Fig. S17**). These results support the finding that six amino acid residues contribute to the compatibility of NRC3 variants to Rpi-blb2, with two K to N changes playing major roles in the NRC3 subfunctionalization process.

**Fig. 4.**
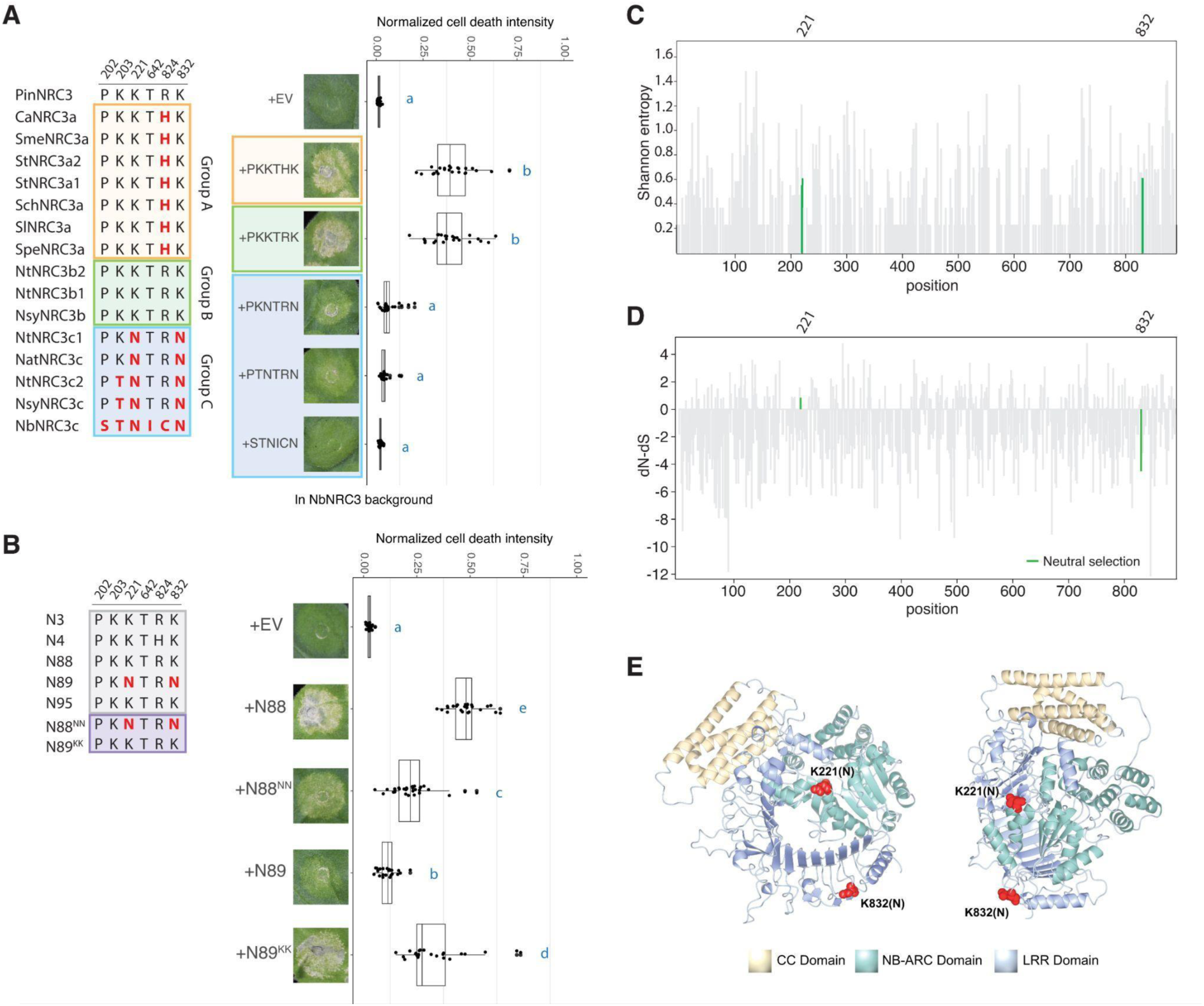
Two K to N mutations play a major role in NRC3 subfunctionalization. (A) Introducing PKKTRK or PKKTHK into NbNRC3c enables it to function with Rpi-blb2. Left panel, the polymorphisms in natural NRC3 variants at the six positions identified. Right panel, cell death assays testing these polymorphisms in NbNRC3c background. (B) Swapping two K and N in N88/N89 changes the compatibility to Rpi-blb2. Left panel, the polymorphisms in ancestral NRC3 variants at the 6 positions identified. Right panel, cell death assay testing two lysine-asparagine changes in both N88 and N89 backgrounds. The NRC3 variants were co-expressed with Rpi-blb2 and AVRblb2 in *nrc2/3/4*_KO *N. benthamiana*. The dot plot represents cell death intensity quantified by UVP ChemStudio PLUS at 6 dpi. Statistical differences were analyzed with Tukey’s HSD test (p<0.05). (C) The entropy analysis of natural NRC3 variants. The protein sequences of NRC3s were aligned using MAFFT and the Shannon entropy was calculated. The positions of the two K to N changes were highlighted in green. (D) dN-dS calculation of natural NRC3 variants by using SLAC analysis. The positions of K to N changes were highlighted in green. The two residues were under neutral selection based on FEL analysis. (E) Predicted protein structure of NbNRC3c in the AlphaFold Protein Structure Database. The CC, NB-ARC, and LRR domains were colored in mist yellow, cyan, and steel blue, respectively. The two positions where K to N changes were highlighted in red.

To understand the polymorphisms of these two positions across NRC3 alleles, we performed entropy analysis using the protein sequence alignment of the natural NRC3 variants. We found that these positions both had entropy values of 0.637 and did not stand out as being among the most conserved or diversified (**Fig. 4C and Data S4**). We then calculated the dN-dS value using SLAC (Single-Likelihood Ancestor Counting) and found that the residue at position 221 showed slightly higher dN than dS values, while the residue at position 832 showed lower dN value than dS value (**Fig. 4D and Data S5**). Despite this, the FEL (Fixed Effects Likelihood) analysis indicated that both positions are under neutral selection (**Fig. 4D and Data S5**). These results suggest that changes in these amino acids among different NRC3 allelic groups that led to subfunctionalization are likely the result of random genetic drift rather than under strong selection. Analysis of these residues on the NbNRC3c structure, predicted by AlphaFold, show that these two amino acids are at positions distant from each other, suggesting that multiple surfaces of the NB-ARC and LRR domains determine the compatibility of helper NLRs to sensor NLRs (**Fig. 4E**).

### Transient interactions between sensor and helper NLRs determine their compatibility

To further dissect the molecular mechanisms of helper-sensor compatibility, we tested the interactions between NRC3 variants with the sensor NLR Rpi-blb2. We focused on two variants, NNN (NbNRC3c) and NN_PKK_N_THK_, as these two variants differ from each other by only six amino acids but display robust differences in their compatibility with Rpi-blb2 (**Fig. 3G**). We co-expressed Rpi-blb2 with NNN or NN_PKK_N_THK_ with or without AVRblb2 and then performed co-immunoprecipitation analyses. When we pulled down Rpi-blb2, we detected very weak signals from NNN and NN_PKK_N_THK_ regardless of whether AVRblb2 was present or not (**Fig. S18A**). Similarly, when we pulled down the two NRC3 variants, we detected weak signals from Rpi-blb2 (**Fig. S18B**). These results suggest that steady-state interactions between sensor and helper NLRs detected using co-IP experiments do not reflect their compatibility, an issue that has indeed limited the use of standard protein interaction assays to study biochemical interactions in the NRC network (*17*, *25*, *26*).

We hypothesized that Rpi-blb2 and NRC3 variants may show differential transient interaction post effector detection, and this transient interaction may be revealed by TurboID-based proximity labeling. To test this, we replaced the constitutive promoter with a copper-inducible promoter to drive the expression of AVRblb2. This chemical inducible system was recently used to regulate the expression of Cas9 for genome editing in *N. benthamiana* (*27*). We found that this inducible system allowed us to precisely activate the cell death responses using copper treatment without any leakiness issue (**Fig. 5A**), and the accumulation of GFP-AVRblb2 was detectable in immunoblot analysis from 3 hours post copper treatment (**Fig. 5B**). We performed proximity labeling by co-expression of NRC3 variants fused to TurboID, RFP-Rpi-blb2, and inducible AVRblb2, and then detected the level of biotinylation on Rpi-blb2 at 6, 12, and 30 hours after copper and biotin treatment (**Fig. 5C**). We found a stronger biotin label signal on Rpi-blb2 at 6 hours post copper/biotin treatment when it was co-expressed with NN_PKK_N_THK_ but not with NNN (**Fig. 5D**). The labeling signal became weaker at 12 hours post-treatment and not detectable at 30 hours post-treatment (**Fig. 5D and Fig. S19**). These results suggested that sensor NLR Rpi-blb2 shows higher transient interaction with the compatible NRC3 variants than the NRC3 variants that are incompatible.

**Fig. 5.**
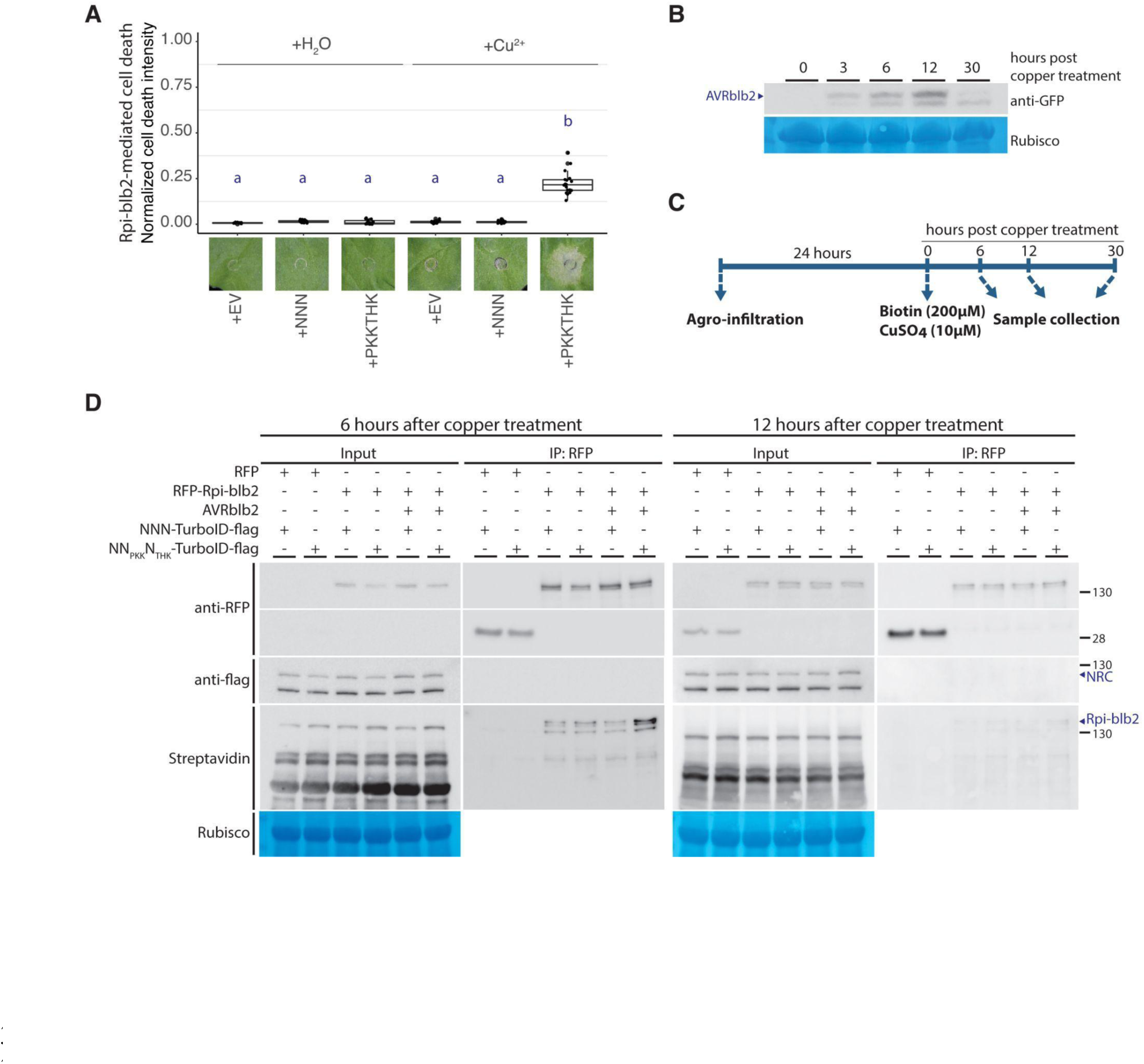
Transient interactions between sensor and helper NLRs determine their compatibility. (A) Quantification of Rpi-blb2-mediated cell death in response to the activation of immunity triggered by copper-inducible flag-AVRblb2 in *nrc2/3/4*_KO *N. benthamiana*. RFP-Rpi-blb2, NRC3s-TurboID-flag (empty vector as a control) were co-expressed with copper inducible flag-AVRblb2 and CUP2-P65. Copper (water as a control) was infiltrated at 1 dpi. The dot plot represents cell death intensity observed at 1-day post copper treatment quantified by UVP ChemStudio PLUS. Statistical differences were analyzed with Tukey’s HSD test (p<0.05). (B) Protein accumulation of copper inducible GFP-AVRblb2 in response to copper treatment. GFP-AVRblb2 was expressed in *nrc2/3/4*_KO *N. benthamiana*. Copper was infiltrated at 1 dpi. Samples were collected at 0, 3, 6, 12, and 30 hours post copper treatment. Western blot was done by using α-GFP antibody. (C) Workflow of TurboID-based proximity labeling assay. Genes-of-interests were transiently co-expressed in *nrc2/3/4*_KO *N. benthamiana* by *Agrobacterium*-mediated transformation. 200 µM biotin and 10 µM CuSO_4_ (or water) were infiltrated into leaves at 24 hpi. Leaf samples were collected at 6, 12, and 30 hours post copper treatment. (D) Western blot of TurboID-based proximity labeling assay of sample collected at 6 and 12 hours post copper/biotin treatment. RFP-Rpi-blb2 and NRC3s-TurboID-flag were co-expressed in *nrc2/3/4*_KO *N. benthamiana*. NRC3s-myc coexpressed with RFP were used as negative controls. All treatments were coexpressed with copper inducible flag-AVRblb2 and CUP2-P65. Treatments with AVRblb2 induction are indicated with ‘+’, while the treatments without AVRblb2 induction are indicated with ‘-’. Protein extracts (input) and RFP-Trap pull-down samples (IP) were analyzed using Western blot analysis using α-RFP, α-flag antibodies, and streptavidine fused with HRP. SimplyBlue SafeStain-staining of Rubisco was used as the loading control.

### Transient interaction between Rpi-blb2 and compatible NRC3 occurs before the formation of membrane-associated puncta

Recent studies suggest that NRCs form membrane-associated punctate upon activation by corresponding sensor NLRs and effectors (*16*, *17*). To test whether the compatibility of NRC3 variants to the sensor NLRs is consistent with punctate formation, we performed cell biology assays by co-expression of NRC3 variants with Rpi-blb2/AVRblb2 or Pto/AvrPto. Co-expressions with Pto/AvrPto triggered both NbNRC3c and SlNRC3a to form membrane-associated punctate, whereas Rpi-blb2/AVRblb2 only triggered SlNRC3a but not NbNRC3c to form membrane-associated punctate (**Fig. 6A-B and Fig. S20**). Consistent with these findings, Rpi-blb2/AVRblb2 can also trigger NRC3 variants NSS, NN_3_N_T5be_, and NN_PKK_N_THK_ to form membrane-associated punctate (**Fig. 6A-B and Fig. S20**). Next, we expressed Rpi-blb2 and NRC3 variants with inducible AVRblb2, and examined the leaves using a confocal microscope at 3, 6, 12, and 30 hours after induction of AVRblb2 expression (**Fig. 6C**). While no punctate was observed at 3 and 6 hours, we observed membrane-associated punctate of NN_PKK_N_THK_ at 12 and 30 hours after AVRblb2 induction (**Fig. 6D-E**). No punctate was observed when NNN or NN_PKK_N_THK_ were expressed with Rpi-blb2 without AVRblb2 expression (**Fig. S21**). These results indicate that transient interaction between Rpi-blb2 and the compatible NRC3 variants occurs before the formation of membrane-associated punctate. Furthermore, NRC3 variants, including SlNRC3a and NN_PKK_N_THKI_, that functioned together with Rpi-blb2 in the cell death assays, also rescued Rpi-blb2-mediated disease resistance in the *nrc2/3/4*_KO *N. benthamiana* (**Fig. 6F**).

**Fig. 6.**
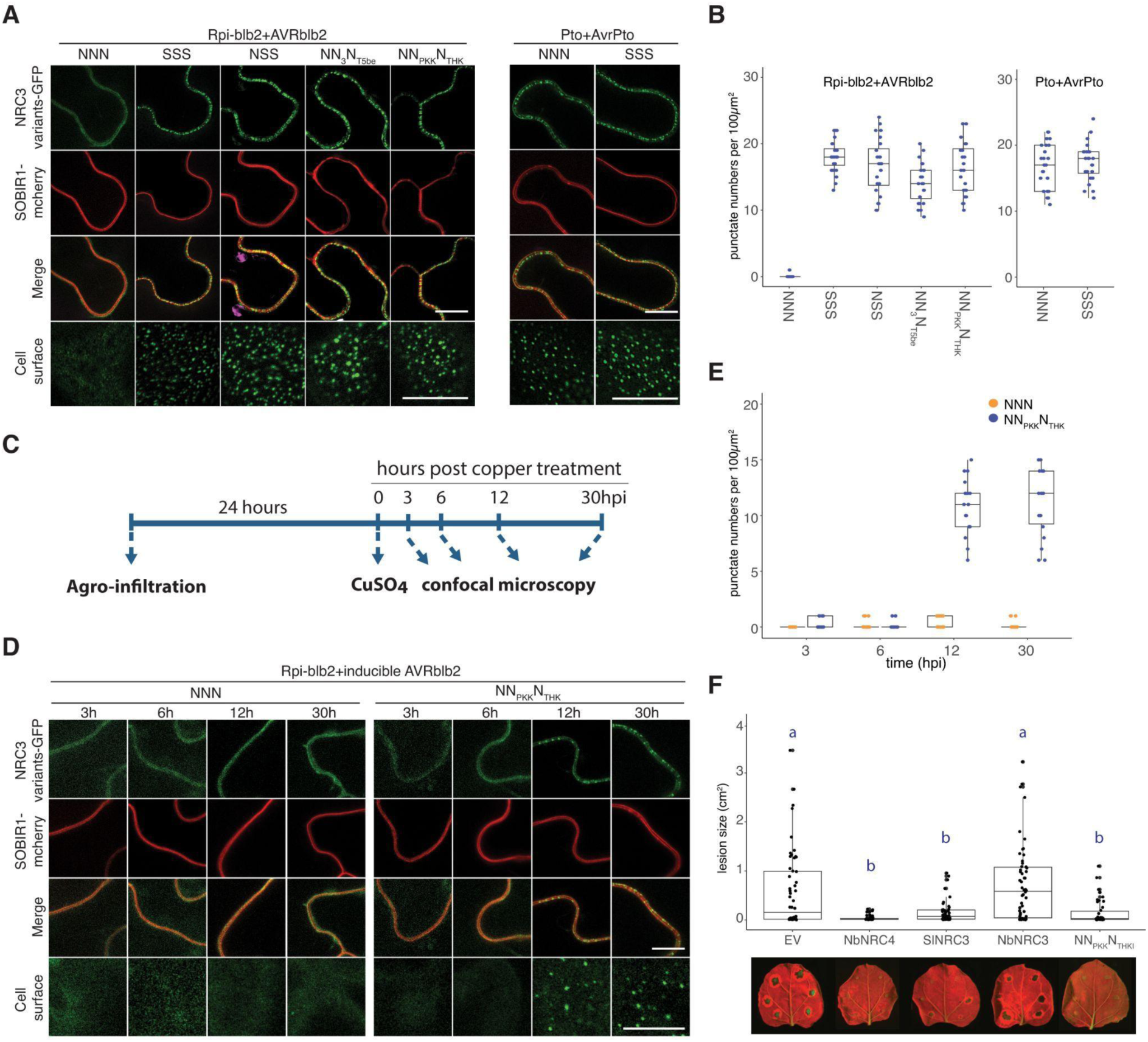
Transient interaction between Rpi-blb2 and compatible NRC3 occurs before the membrane-associated puncta formation. (A) Compatible NRC3 variants form membrane-associated punctate upon immune activation. NRC3 variants fused with GFP were co-expressed with Rpi-blb2/AVRblb2 or Pto/AvrPto. Samples were examined at 3 dpi. Scale bars represent 10µm. SOBIR1-mcherry was used as a plasma membrane marker. (B) Quantification of punctate of NRC3 variants in (A). (C) Workflow for detection of punctate formation after induction of AVRblb2 expression. Genes-of-interests were transiently co-expressed in *nrc2/3/4*_KO *N. benthamiana* by *Agrobacterium*-mediated transformation. Water or 10 µM CuSO_4_ were infiltrated into leaves at 24 hpi. Leaf samples were examined at 6, 12, and 30 hours post copper treatment. (D) NN_PKK_N_THK_ forms membrane-associated punctuate dots in response to the activation of immunity triggered by copper-inducible flag-AVRblb2. Hellfire-tagged Rpi-blb2 and NRC3 variants fused with GFP were co-expressed with copper-inducible flag-AVRblb2 and CUP2-P65. Copper was infiltrated at 24 hpi. Samples were examined at 3, 6, 12, and 30 hours post copper treatment. Scale bars represent 10µm. SOBIR1-mcherry was used as a plasma membrane marker. (E) Quantification of punctate dots of NNN and NN_PKK_N_THK_ in 3, 6, 12, and 30 hours post copper treatment. (F) Infection assay of *P. infestans* in *nrc2/3/4*_KO *N. benthamiana* transiently expressing Rpi-blb2 and NRC variants. The dot plots represent the lesion sizes observed at 5 days post-infection quantified by UVP ChemStudio PLUS. Statistical differences were analyzed with Tukey’s HSD test (p<0.05).

## Discussion

The NRC network is crucial for pathogen resistance in solanaceous crops, but the determinants and evolution of its complex structure have been elusive. Here, by exploring ancestral and natural variants of NRC homologs, we reveal that subfunctionalization is key to the evolution of the NRC network. By examining closely related NRC3 orthologs, we identified critical polymorphisms in helper-sensor compatibility, pinpointing three amino acids in both the NB-ARC and LRR domains. Notably, two lysine-to-asparagine mutations significantly impact helper-sensor compatibility and subfunctionalization. These specific amino acid differences influence the transient interaction between helper and sensor NLRs upon pathogen effector detection. Our findings thus unravel both mechanistic and evolutionary aspects underpinning the complex genetic architecture of the NRC network.

How did subfunctionalization events impact the evolution of NLR networks? We propose that the subfunctionalization of NRCs led to the division of the NLR network into smaller subnetworks (**Fig. 7**). However, evolutionary analyses indicated that residues that affect NRC3 compatibility to Rpi-blb2 are mostly under neutral selection, suggesting that subfunctionalization may be the result of random genetic drift accumulating over time rather than being due to strong selection. The evolution from ancestral NRC helper-sensor gene cluster to the complex network with partially redundant NRC2/3/4 nodes in solanaceous plants may have been the result of successive cycles of gene duplications followed by subfunctionalizations (**Fig. 7**). Interestingly, NRC networks are highly expanded independently in several lamiids, with many NRCs showing partial redundancy, such as in both Solanaceae and Convolvulaceae (*13*, *14*). NRC networks with intricate genetic architectures must have provided some long-term advantages for the plant species to survive. One possibility is that this helped improve the robustness of the immune system, becoming more resistant to pathogen perturbation. Such examples include SPRYSEC15 and AVRcap1b effectors that target NRC2 and NRC3, but not NRC4 (*28*). If little or no diversification and subfunctionalization occurs among NRC homologs, a single or few effectors that suppress the function of NRCs could have dramatically compromised the NRC networks.

**Fig. 7.**
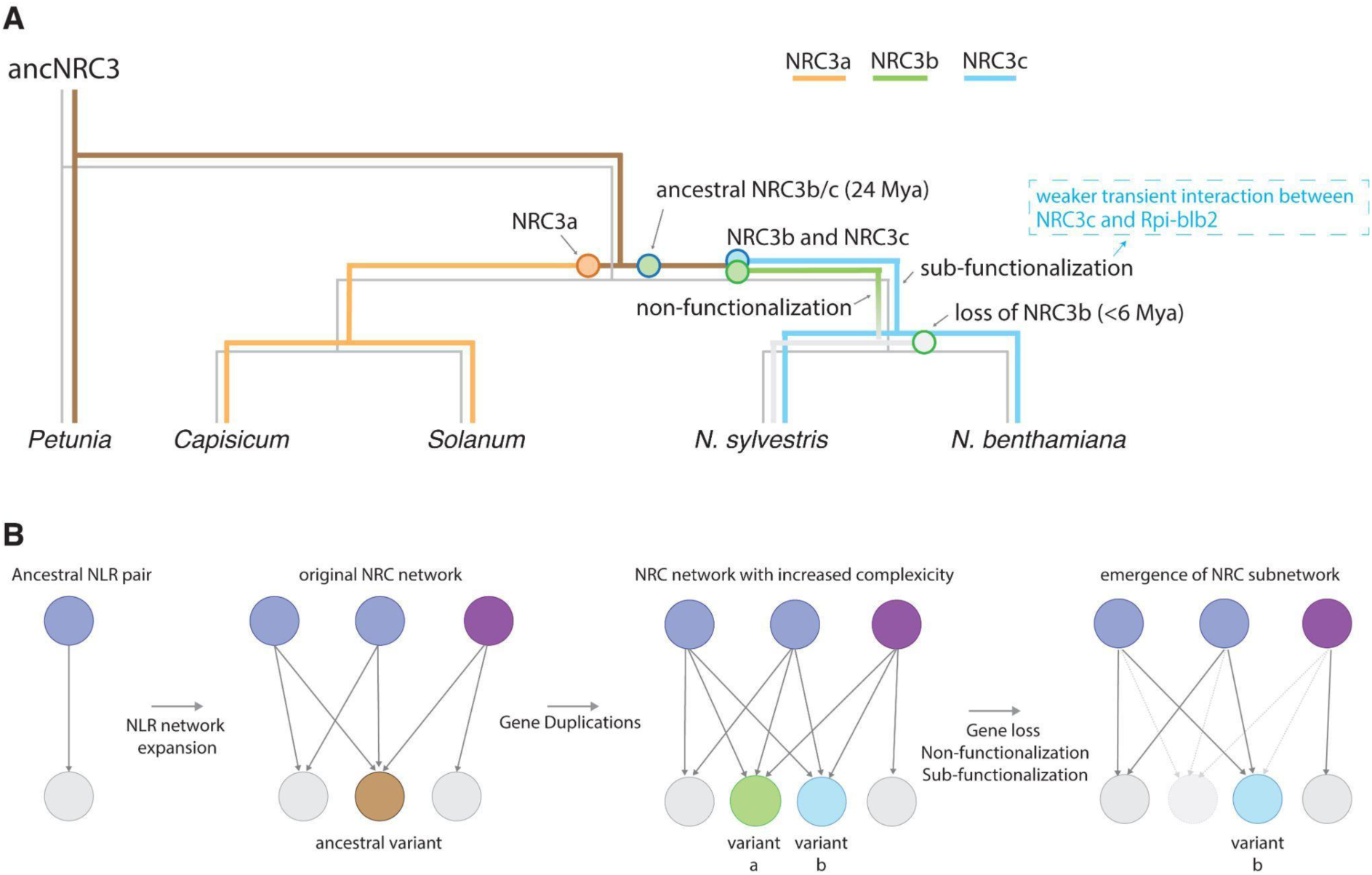
Functional divergence and gene loss of NRCs led to the division of the NLR network into smaller subnetworks. (A) Evolutionary history of NRC3. The ancestral NRC3 (ancNRC3) first duplicated into two alleles, NRC3a in *Capsicum* and *Solanum* species and ancestral NRC3b/c in *Nicotiana* species. The ancestral NRC3b/c was further duplicated to generate NRC3b and NRC3c. NRC3b later underwent nonfunctionalization and was lost in *N. benthamiana* while NRC3c underwent subfunctionalization that led to weaker transient interaction between Rpi-blb2 and NRC3c. As a result, the NRC3 in *Nicotiana* species lost the ability to function with Rpi-blb2. (B) Development of the NRC network in solanaceous plants. The original NLR pair expanded to form an initial NRC network with an ancestral NRC (brown), functioning with multiple sensor NLRs. This ancestral variant duplicated, resulting in two functionally redundant variants: ’a’ and ’b’ (green and blue, respectively). Variant ’a’ lost its sensor NLR compatibility due to gene loss or nonfunctionalization, while variant ’b’ retained partial compatibility, undergoing subfunctionalization. Consequently, variant ’b’ now functions with a narrower range of sensor NLRs, forming a distinct NRC subnetwork.

The difference in signaling compatibilities between orthologous helper and sensor NLRs has been observed in many other cases. Since the NRC network originated from an ancestral NLR pair, the phenomenon observed in paired NLRs may resemble the early scenario in NRC network evolution. In Arabidopsis, two orthologous RPS4/RRS1 NLR pairs were reported. Members in these two pairs, RPS4A/RRS1A and RPS4B/RRS1B, only function with their authentic partners (*29*). While the TIR domain is indispensable for the specificity between RPS4A/RRS1A and RPS4B/RRS1B pairs, swapping the NB-ARC, LRR, or DOM4 of RRS1B into RRS1A all compromised the cell death responses when co-expressed with RPS4A and the effector (*30*). In rice, the helper NLR Pias-1 can also function with the sensor NLR RGA5 to respond to AVR-Pia, indicating that Pias-1 is not specialized to its own linked sensor NLR Pias-2 (*31*). In contrast, helper-sensor specialization was observed in allelic Pik-1/Pik-2 rice NLR pairs (*32*). For Pikm-1/Pikm-2 and Pikp-1/Pikp-2, this helper-sensor specialization was mapped to a single amino acid polymorphism on Pik-2 that determines its preferential association between matching pairs (*33*). Presuming that orthologous paired NLRs should exhibit cross-compatibility immediately following the initial duplication, various degrees of subfunctionalization have occurred throughout the co-evolution of distinct paired NLRs and their orthologous counterparts.

Our research primarily examined mutations in helper NLRs and their influence on compatibility with sensor NLRs. Given that specific amino acids are integral to transient interactions between these NLR types, it is plausible that sensor NLRs also possess corresponding residues dictating their compatibility with helper NLRs. These residues might determine the sensor NLRs’ ability to signal through a broader or narrower array of NRCs. Notably, the sensor NLR Rx is distinguished by its capacity to signal through a wider range of NRCs compared to other tested sensor NLRs, functioning with solanaceous NRC2/3/4 and multiple NRCs in *Ipomoea aquatica* (*14*). Recent findings suggest that the NB domains of various sensor NLRs are sufficient to induce NRC oligomerization and trigger immune signaling, highlighting the NB-ARC domain’s key role in sensor-helper compatibility from the sensor perspective (*34*). Investigating additional natural variants and analyzing the residues involved in helper-sensor NLR compatibility across different sensor and helper NLRs could further elucidate how the NLR network’s signaling connections maintain their specificity and promiscuity. Finally, another intriguing aspect yet to be fully explored is the signal transduction mechanism from sensor to helper NLRs. The activation-and-release model has been proposed to explain helper NLR activation by sensor NLRs following pathogen effector detection (*17*, *25*). Our findings indicate that transient interactions between sensor and helper NLRs occur during immune activation. Capturing and resolving the structures of these transient helper-sensor NLR complexes could shed light on the intricate mechanisms governing helper-sensor compatibility and signal transduction.

## Materials and Methods

### Plant material and growth conditions

Wild type (WT) and *nrc2/3/4*_KO *N. benthamiana* (*21*) were grown in a walk-in chamber with a temperature of 25°C, humidity of 45–65%, and 16/8 hr light/dark cycle.

### *Phytophthora infestans* and disease resistance assay

*Phytophthora infestans* 214009 (a gift from Dr. Jin-Hsing Huang, Taiwan Agriculture Research Institute) was isolated from Taiping, Taichung City, Taiwan. *P. infestans* was maintained in the Rye medium at 19°C in dark conditions (*35*). Before the infection assay, the *P. infestans* was subcultured on tomato (money maker) leaf discs. In the disease resistance assay, four-week-old N. benthami*ana* plants were vacuum infiltrated with *A. tumefaciens* containing constructs of interest shown in Table S2. Sporangia were collected from *P. infestans*-infected tomato leaf discs by gently shaking at 4°C sterile water. Droplets of 10μL of sporangia suspensions (3 x 10^3^ sporangia/mL) were applied to the abaxial side of detached *N. benthamiana* leaves 6 hours post vacuum infiltration. The infected leaves were placed in the Corning® 245 mm Square Dish at 19°C in dark condition. The disease symptoms were quantified using UVP ChemStudio Imaging Systems at 5 days post-infection (dpi). Blue LED light was used for excitation and FITC filter (519 nm) and Cy5 filter (670 nm) were used to detect the autofluorescence from infected and healthy tissues.

### Plasmid constructions

All the NRC3 variants used in this study were constructed using Golden Gate assembly (*36*, *37*). The natural variants were amplified from genomic DNA and cloned into a binary vector (pICH86988) which carries the 35S promoter and OCS terminator. The petunia NRC3 was synthesized and then cloned into pICH86988. The auto-active mutants of N88, N89, and NRC3bs were generated through site-directed mutagenesis introducing a D to V mutation in the MHD motif. The chimeric NRC3 variants were made by modularizing regions of NbNRC3 and SlNRC3 as level 0 modules and re-assembled by using Golden Gate assembly. The CC, NB-ARC, and LRR domains were first modularized as level 0 modules to investigate which domain contributes to the subfunctionalization of NbNRC3c. The stop codon in the LRR domains was removed for tagging with C-terminal tags. These modules were assembled into pICH86988 together with C-terminal 4x myc. To identify the residues that contribute to the subfunctionalization in the NB-ARC domain, the polymorphisms in this domain were introduced into the NbNRC3 NB-ARC module to generate 10 NB-ARC modules. These modules were assembled with CC and LRR domains to obtain full-length NRC3 variants. The polymorphisms in the LRR domain were analyzed using a similar strategy. We first divided this domain into 5 fragments (1 to 5) and made them into independent level 0 modules. We found that the polymorphisms in regions 2 and 5 corporately contributed to the subfunctionalization. Therefore, we further mutated LRRmodules of NbNRC3 to introduce the polymorphisms into these regions. These modules were assembled with CC, NB-ARC, and other fragmented LRR modules to become full-length NRC3 variants. All modules, together with C-terminal 4x myc, were assembled into pICH86988. The lysine-asparagine swapping in the N88 and N89 background was done by site-directed mutagenesis. For the inducible system, AVRblb2 was placed under the copper-inducible promoter (CBS4), while the DNA-binding domain fused with the activation domain (CUP2-VP16) was placed under the 35S promoter. Sequences of plasmids, NRC3 variants, and primers used in this study are listed in Data S1-3.

### Agroinfiltration

We used *Agrobacterium tumefaciens* to transiently express the NRC3 variants, sensor NLRs, and effectors in four-week-old *N. benthamiana*. Strains of *A. tumefaciens* were refreshed from glycerol stock to 523 medium containing appropriate antibiotics at 28 °C overnight. Cells were harvested by centrifugation at 2500 × g, room temperature for 5 min. Cells were resuspended in MMA buffer (10 mM MgCl2, 10 mM MES-KOH, 150 µM acetosyringone, pH5.6) to the concentration listed in Table S1 and then infiltrated into leaves using 1 mL syringes.

### Cell death assay

The complementation assays were conducted in *nrc2/3/4*_KO *N. benthamiana* background, while auto-activity analyses were done in WT *N. benthamiana*. Sensor NLRs and corresponding effectors were transiently expressed by *A. tumefaciens* strains containing expression vectors for different proteins as indicated. The cell death intensity was quantified using UVP ChemStudio Imaging Systems at 6 days post agroinfiltration (dpi). We set blue light as excitation and used a FITC filter (519 nm) to detect the autofluorescence of necrotic regions. Cell death intensity was normalized by dividing the pixel value by the maximum pixel value of the signal (65535).

### Protein extraction

Leaf tissue was finely powdered by grinding in a mortar and pestle with liquid nitrogen and total protein was subsequently extracted with extraction buffer (10% glycerol, 25 mM Tris pH7.5, 1 mM EDTA, 150 mM NaCl, 2% w/v PVPP, 10 mM DTT, 1x protease inhibitor cocktail (Sigma), 0.1% IGEPAL (Sigma)). After centrifugation at 13,000 g for 10 min at 4°C, the supernatant fraction was taken out and mixed with 4x sample loading dye (200 mM Tris-HCl (PH6.8), 8% (w/v) SDS, 40% (v/v) glycerol, 50 mM EDTA, 0.08% bromophenol blue) with 100 mM DTT. Protein samples were incubated at 70°C for 10 minutes before being analyzed by SDS-PAGE.

### SDS-PAGE electrophoresis and immunodetection analyses

SDS-PAGE electrophoresis and immunodetection analyses were conducted as previously described (*38*, *39*). Denatured samples were run on PAGE containing 10% or 15% T-Pro EZ Gel Solution, 0.1% (v/v) TEMED (Bio-Rad), and 0.1% (w/v) ammonium persulfate (Bio-Rad). The proteins were then transferred to PVDF membranes using the Trans-Blot Turbo Transfer System (Bio-Rad). Immunoblotting and detection were performed on the PVDF membrane with SNAP i.d.® 2.0 Protein Detection System (Merck) by using anti-myc (A00704, GenScript), anti-RFP (YH80520, Yao-Hong), anti-flag (F1804, Sigma) and anti-GFP (A-11122, Invitrogen) as primary antibodies and Peroxidase AffiniPure Goat Anti-Mouse IgG (H+L) (115-035-003, Jackson) or Peroxidase Conjugated Goat Anti-Rabbit IgG (H+L) (AP132P, Sigma) as secondary antibodies. First antibodies were at a dilution of 1:8000 while secondary antibodies were at a dilution of 1:25000. The chemifluorescence was detected using SuperSignal West Pico PLUS Chemiluminescent Substrate (34580, Thermo Scientific) mixed with SuperSignal West Femto Maximum Sensitivity Substrate (34096, Thermo Scientific, ratio=4:1). The images were taken under UVP ChemStudio Imaging Systems. SimplyBlue SafeStain (465034, Invitrogen) was applied to detect the signal of rubisco on the PVDF membrane.

### Detection of NRCs protein accumulation

To detect the protein accumulation of NRC3 variants, four-week-old *N. benthamiana* plants were infiltrated with *A. tumefaciens* containing constructs of NRC3 variants-myc at OD_600_ 0.5. Six leaf discs (1.1 cm diameter) were collected at 2 dpi and protein accumulation was detected according to the immunoblot analysis described above.

### Co-immunoprecipitation assay

To detect the steady-state interaction between Rpi-blb2 and NRC3s, four-week-old *N. benthamiana* plants were infiltrated with *A. tumefaciens* containing constructs of interest shown in Table S3. Six leaves (2.5g) were collected at 30 hpi and extracted with the extraction buffer described above. Anti-c-myc beads (VF299569, Thermo) or RFP-Trap Magnetic Agarose (Chromotek) were added to the protein extracts and incubated at 4°C for 3 h. After incubation, the protein mixtures were washed with wash buffer (10% glycerol, 25 mM Tris pH7.5, 1 mM EDTA, 150 mM NaCl, and 0.1% IGEPAL) five times to remove non-specific binding proteins. To elute proteins, beads or Magnetic Agarose were mixed with 1x sample loading dye with 100 mM DTT and incubated at 70°C for 10 minutes before analyzing by SDS-PAGE. Protein signals were detected through the process described above.

### TurboID-based proximity labeling assay

To detect the transient interaction between Rpi-blb2 and NRC3s, four-week-old *N. benthamiana* plants were infiltrated with *A. tumefaciens* containing constructs of interest shown in Table S4. 10 µM CuSO_4_ (or water as a control) was co-infiltrated with Biotin (200 µM) (Sigma) at 24 hpi. Six leaves (2.5g) were collected at 0, 3, 6, and 12 hours after copper (or water) treatment. RFP-Trap Magnetic Agarose was added to the protein extracts and incubated at 4°C for 3 h. After incubation, the protein mixtures were washed with wash buffer for five times to remove non-specific binding proteins. To elute proteins, Magnetic Agarose was mixed with 1x sample loading dye with 100 mM DTT and incubated at 70°C for 10 minutes before analyzing by SDS-PAGE. HRP-Conjugated Streptavidin (VJ306909, Thermo) was used for the detection of biotinylated proteins. Protein signals were detected using the immunodetection process described above.

### Entropy Analysis and Protein Structure Prediction

The sequences of NRC3 of tomato (*Solanum pennellii*, *Solanum chilense*, and *Solanum lycopersicum*), tobacco (*Nicotiana tabacum* and *Nicotiana benthamiana*), pepper (*Capsicum annuum*), eggplant (*Solanum melongena*), and potato (*Solanum tuberosum*) were downloaded from Sol Genomics Network (SGN). The sequences of NRC3 of *Nicotiana sylvestris* were downloaded from NCBI. The accession numbers are listed in Data S3. The protein sequences of NRC3s were aligned using MAFFT (version 7) and used for the entropy analysis. The Shannon entropy was calculated using an online tool established by Los Alamos National Laboratory (https://www.hiv.lanl.gov/content/sequence/HIV/HIVTools.html). A score of 1.5 was set as a cutoff to determine highly variable residues (*40*). Detailed results of the entropy analysis are included in Data S4. The structure of NbNRC3c was predicted by the AlphaFold Protein Structure Database.

### Phylogenetic analysis and detection of selection

NRCs from *Capsicum annuum*, *Solanum lycopersicum*, *Nicotiana benthamiana*, *and Solanum tuberosum* were downloaded from SGN. The full-length nucleotide sequences of NRC or NRC3 variants were translated into protein sequences and aligned by using MEGA software (Molecular Evolutionary Genetics Analysis). The gaps within alignments were removed manually. The trimmed amino acid sequences were applied to phylogenetic analysis by using Maximum-likelihood phylogenetic analysis with the evolutionary model JTT+G+I and 100 bootstrap tests. The aligned NRC3 protein sequences were reversed-translated into nucleotide sequences for detection of selection. The sequences were analyzed by SLAC (Single-Likelihood Ancestor Counting) analysis and FEL (Fixed Effects Likelihood) analysis on Datamonkey Adaptive Evolution Server (https://www.datamonkey.org/) (*41–43*). Detailed results of the SLAC and FEL analyses are included in Data S5.

### Cell biology assays

To detect the punctate formation of NRCs in response to immune activation, four-week-old *N. benthamiana* were agroinfiltrated with constructs of interest shown in Table S5. Leaf tissues were imaged at 3 dpi by using Olympus FV3000 with a 60x silicone oil immersion objective with the excitation of 488 nm for GFP and 561 nm for mCherry. To dissect the timepoint of punctate dots formation of NRC3s after AVRblb2 induction, four-week-old *N. benthamiana* were agroinfiltrated with constructs of interest shown in Table S5. Leaf tissues were imaged at 3, 6, 12, and 30 hours post copper (or water) treatment.

### Ancestral sequence reconstruction

Sequences of the NRC family members were extracted from 124 Solanaceae genomes using NLRtracker followed by a phylogeny-based approach (detailed in the Zenodo repository). To identify the NRCX/1/2/3 clades, we aligned the 1,116 protein sequences of the NB-ARC domain of the NRC family using MAFFT v7.508 with default options. Sequences of NB-ARC regions extracted in Adachi et al., 2023 were also included in the analysis (*23*). We then reconstructed a phylogenetic tree using FastTree v2.1.11 (*44*) and extracted 324 NRC sequences within the NRCX/1/2/3 clades. To obtain a codon-based nucleotide sequence alignment, we re-aligned 324 full-length protein sequences of the NRCX/1/2/3 clades using MAFFT with default options and, then, we threaded the nucleotide sequences onto the protein alignment using the “thread_dna” command in phykit v1.11.14 (*45*). Based on this nucleotide sequence alignment, we reconstructed the phylogenetic tree of NRCX/1/2/3 using IQ-TREE v2.2.0.3 (http://www.iqtree.org) (*46*) with 1,000 ultrafast bootstrap replicates (*47*). The best-fit model to reconstruct the tree was automatically selected by ModelFinder (*48*) in IQ-TREE, and the “TIM3+F+I+I+R4” model has been selected according to the Bayesian Information Criterion (BIC). Based on this phylogenetic tree and codon-based alignment, we reconstructed ancestral sequences using FastML with the nucleotide sequence mode and its default settings (Data S6). The five selected NRC3 ancestral variants were synthesized using the service provided by SynBio Technologies (New Jersey, USA) and then cloned into pICH86988.

## Acknowledgments

We thank Dr. Sophien Kamoun (The Sainsbury Laboratory, United Kingdom) for suggestions about the experiments and manuscript. We thank Mark Youles (SynBio, The Sainsbury Laboratory, UK) for sharing plasmids for molecular cloning. We thank Mei-Jane Fang and Ji-Ying Huang in the Live-Cell-Imaging Core Lab (Institute of Plant and Microbial Biology, Academia Sinica, Taipei, Taiwan) for help with confocal imaging. We thank Dr. Jin-Hsing Huang (Taiwan Agriculture Research Institute) for providing the *P. infestans* isolated 214009 and advice on pathogen inoculation.

## Funding

National Science and Technology Council NSTC-110-2311-B-001-044 (CYH, YSH, JCLA, CHW)

National Science and Technology Council NSTC-111-2628-B-001-023 (CYH, YSH, HYW, YFC, BJC, CHW)

National Science and Technology Council NSTC-112-2628-B-001-007 (HYW, YFC, BJC, CHW)

National Science and Technology Council NSTC-112-2813-C-001-025-B (KYL, CHW)

Institute of Plant and Microbial Biology, Academia Sinica, intramural fund (CYH, YSH, LTH, JCLA, CHW)

National Institute of Agricultural Botany (NIAB) Fellowship (LD) The Gatsby Charitable Foundation (YS, AT, JK, LD)

The Royal Society (LD) BASF Plant Science (JK)

## Author contributions

Conceptualization: CYH, YSH, LD, CHW

Data curation: CYH, YSH, YS, LTH, AT, JK, CHW

Formal analysis: CYH, YSH, CHW

Investigation: CYH, YSH, LD, CHW

Methodology: CYH, YSH, YS, HYW, YFC, KYL, BJC, CHW

Resources: CYH, YSH, JCLA, CHW

Supervision: CHW

Funding acquisition: CHW

Project administration: CHW

Writing initial draft: CYH, YSH, LD, CHW

Editing: CYH, YSH, YS, JCLA, YFC, JK, LD, CHW

## Data and materials availability

All data are available in the main text or the supplementary materials. Datasets associated with ancestral sequence reconstruction are available on Zenodo (https://doi.org/10.5281/zenodo.10354350; https://doi.org/10.5281/zenodo.10360199)

