## Supplemental Figs S1-S21, Tables S1-S5 for "Functional divergence shaped the network architecture of plant immune receptors"

**This PDF file includes:**

Figs. S1 to S21
Tables S1 to S5
References (1 to 13)

**Other Supplementary Materials for this manuscript include the following:**

Data S1 to S6

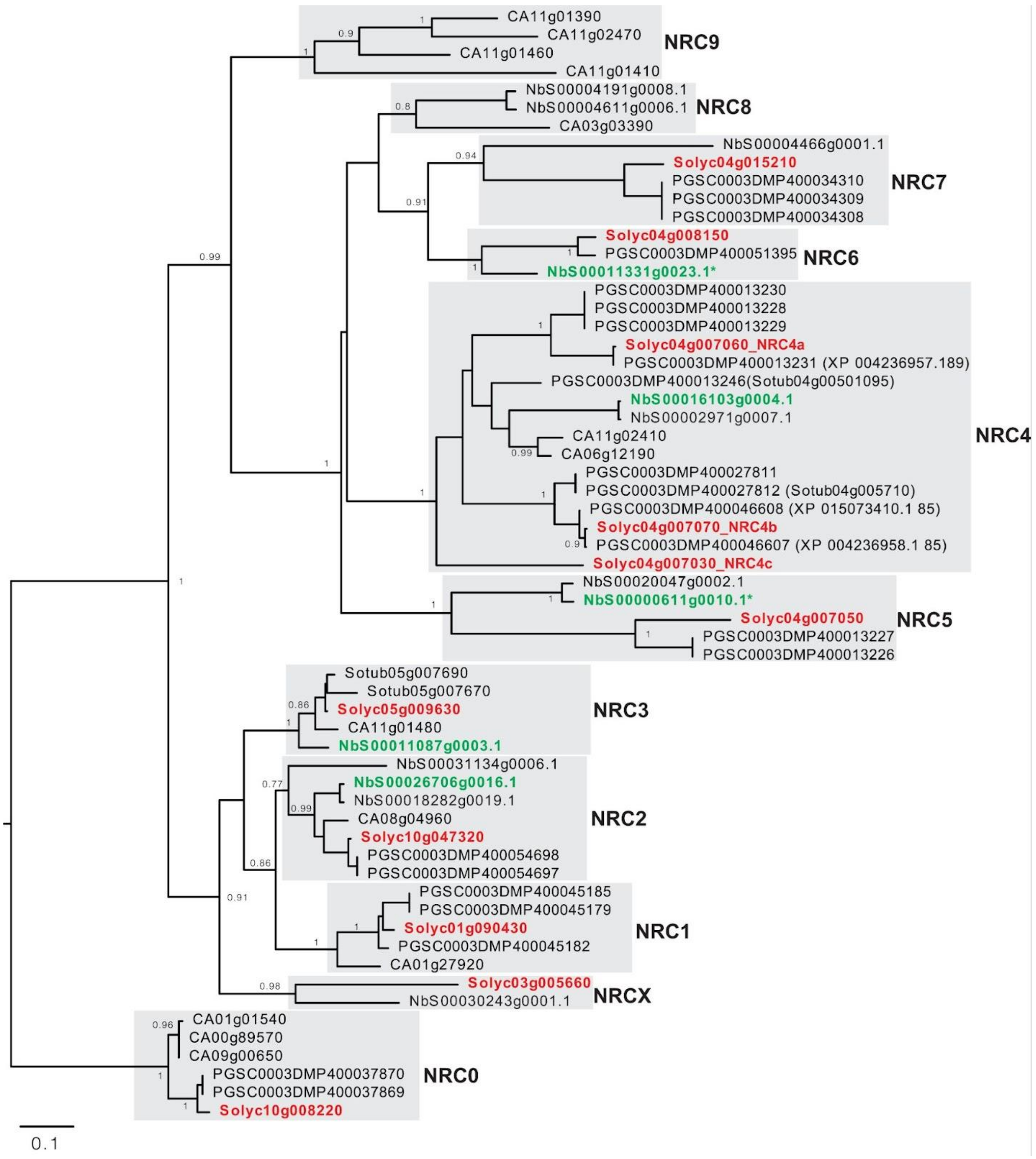


**Fig. S1.**

The NRC from solanaceous plants were grouped into NRC0 to NRCX based on the phylogenetic analysis. Phylogenetic tree of NRC from tomato, tobacco, potato, and pepper. The sequences of NRCs were downloaded from the Sol Genomics Network. The alignment of amino acid sequences of the NB-ARC domain was used for phylogenetic analysis using the Maximum-likelihood method with 100 bootstrap tests. Gray boxes indicated different NRC clades. The clade of NRC0 was used as an outgroup. Tomato and *N. benthamiana* NRCs used in Fig. 1A and Fig. S2 were highlighted in red and green, respectively.

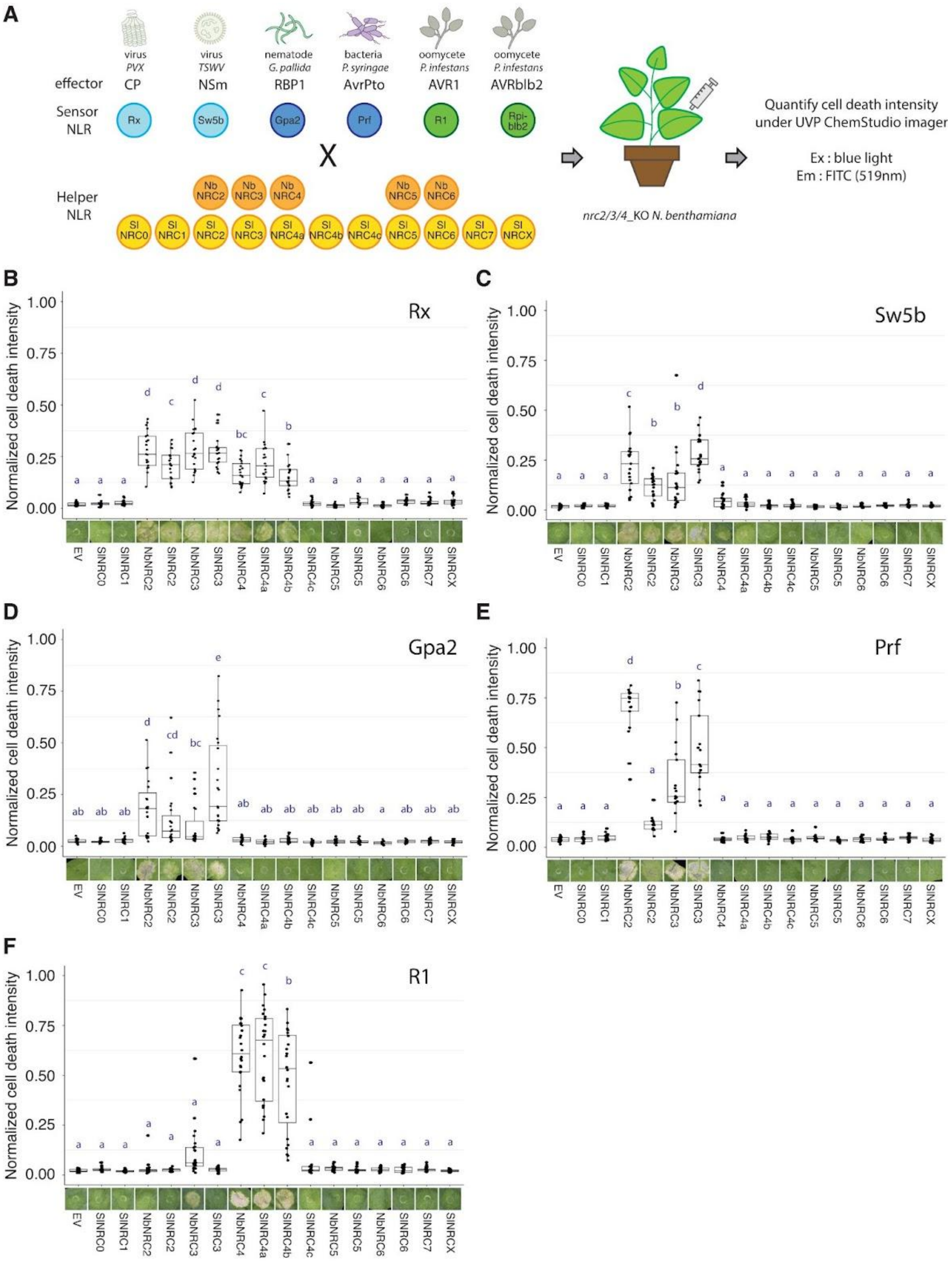
A
virus
PVX
virus
TSWV
nematode
G. pallida
bacteria
P. syringae
oomycete
P. infestans
oomycete
P. infestans
effector
CP
NSm
RBP1
AvrPto
AVR1
AVRblb2
Sensor NLR
Rx
Sw5b
Gpa2
Prf
R1
Rpi-blb2
X
Helper NLR
Nb NRC2
Nb NRC3
Nb NRC4
Nb NRC5
Nb NRC6
Sl NRC0
Sl NRC1
Sl NRC2
Sl NRC3
Sl NRC4a
Sl NRC4b
Sl NRC4c
Sl NRC5
Sl NRC6
Sl NRC7
Sl NRCX
nrc2/3/4_KO N. benthamiana
Quantify cell death intensity under UVP ChemStudio imager
Ex : blue light
Em : FITC (519nm)
B
Rx
C
Sw5b
D
Gpa2
E
Prf
F
R1
Normalized cell death intensity
1.00
0.75
0.50
0.25
0.00
EV
SlNRC0
SlNRC1
NbNRC2
SlNRC2
NbNRC3
SlNRC3
NbNRC4
SlNRC4a
SlNRC4b
SlNRC4c
NbNRC5
SlNRC5
NbNRC6
SlNRC6
SlNRC7
SlNRCX

**Fig. S2.**

NRCs from solanaceous plants displayed diverse compatibility with sensor NLRs. (A) Workflow of cell death assays testing the ability of NRC to function with different sensor NLRs. NRCs were co-expressed with sensor NLRs and the corresponding effectors in *nrc2/3/4*_KO *N. benthamiana*. NRC variants were co-expressed with (B) Rx/CP, (C) Sw5b/NSm, (D) Gpa2/RBP1, (E) Pto/AvrPto, or (F) R1/AVR1 to analyze their ability to work with sensor NLRs. Cell death phenotypes were recorded at 6 dpi. The dot plots represent cell death intensity quantified using UVP ChemStudio PLUS. Statistical differences were analyzed with Tukey's HSD test ($p<0.05$).

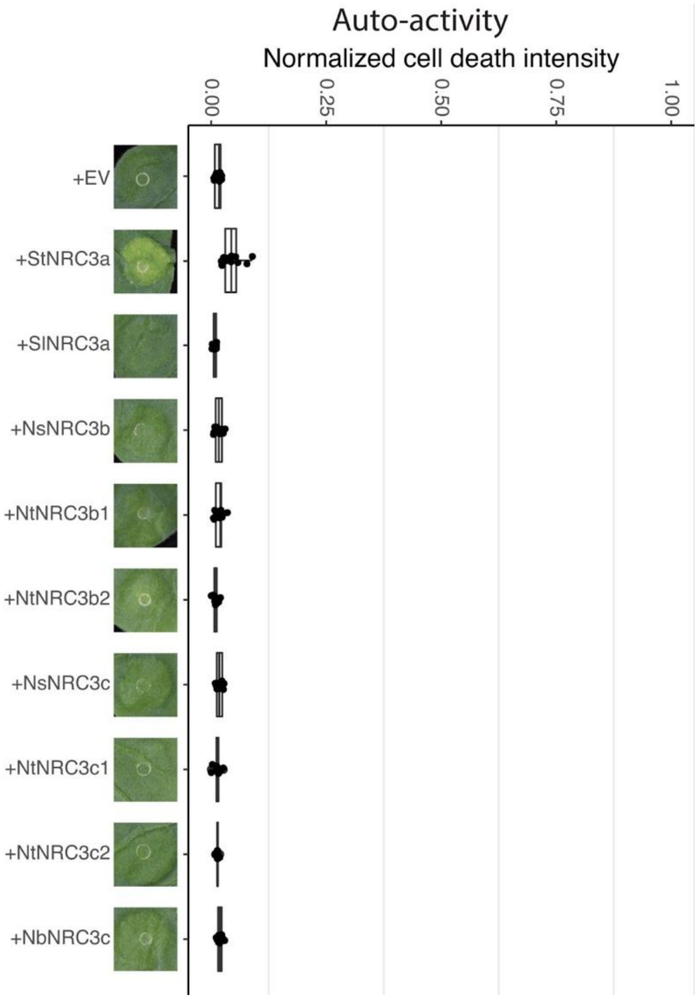


**Fig. S3.**

NRC3 natural variants show no or low auto-activities when expressed in *N. benthamiana*. Auto-activity analysis of NRC3 natural variants tested in Fig. 1D. The NRC3 variants were expressed alone in WT *N. benthamiana.* The dot plots represent cell death intensity quantified using UVP ChemStudio PLUS at 6 dpi.

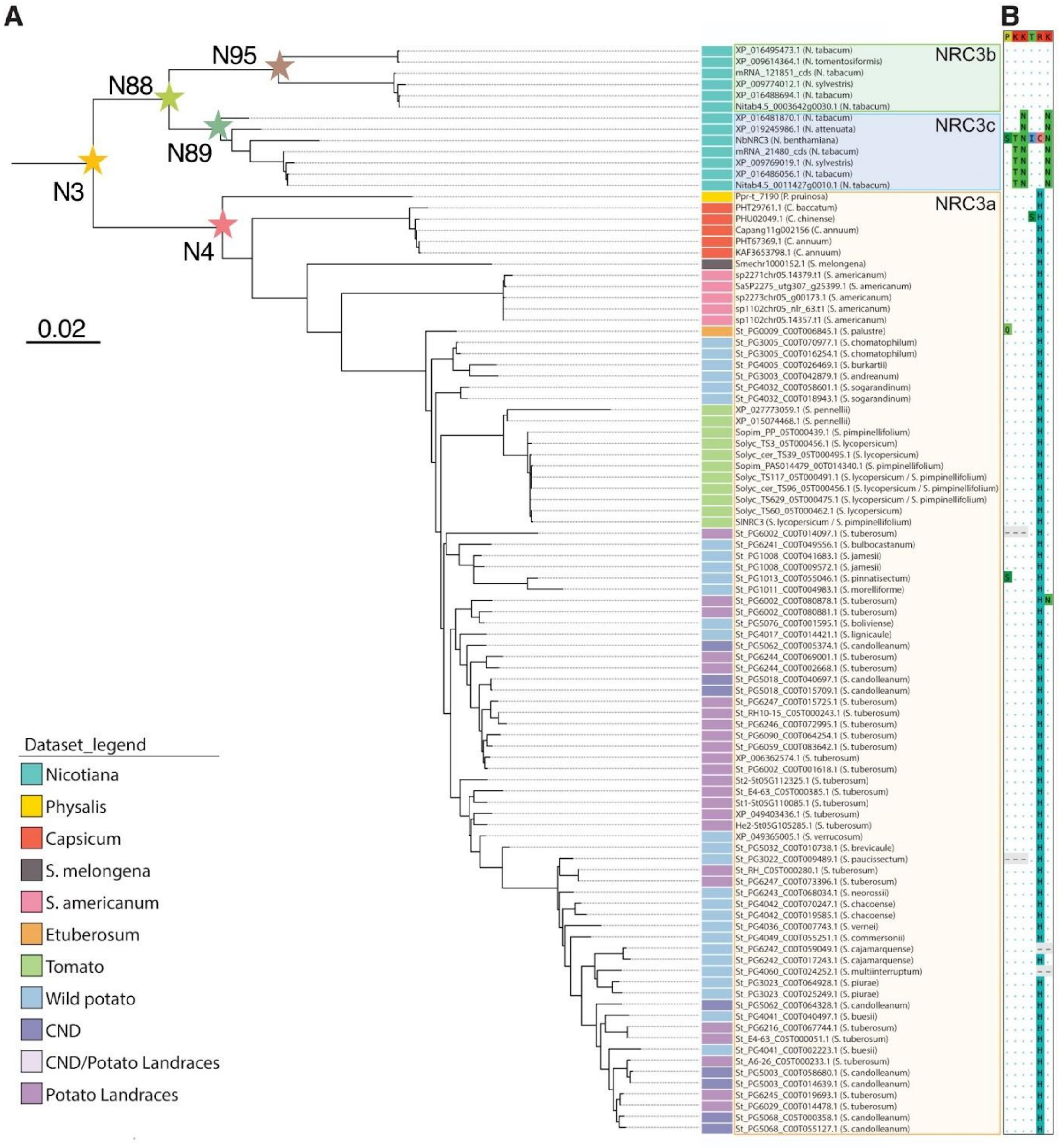


**Fig. S4.**

Ancestral sequence reconstruction of NRC3. (A) Phylogenetic tree of the solanaceous NRC3 clade. The NLR sequences were extracted from 124 genomes in the Solanaceae family using the NLRtracker software. The phylogenetic tree was reconstructed based on the codon-based nucleotide sequence alignment of NRCX, NRC1, NRC2, and NRC3 clades using IQ-TREE. Ancestral sequence reconstruction was performed using FastML. Orange, green, and blue boxes indicate the allelic groups of NRC3a, NRC3b, and NRC3c, respectively. The nodes of ancestral variants tested in this study were indicated with star shapes. N3: the ancestral variant before the divergence of the three allelic groups; N4: the ancestral variant of NRC3a; N88: the ancestral variant before the divergence of NRC3b/c; N95: the ancestral variant of NRC3b; N89: the ancestral variant of NRC3c. (B) Polymorphisms at the six positions identified across NRC3 natural variants. The ancestral states of the six positions at N3 were PKKTRK as listed at the top. Polymorphisms carried by NRC3 variants  are highlighted while the conserved residues are shown as dots.

```
N3    MADVAVKFLVENLMQLLIDNADLILGIKGEVENLLQDLNDFNAFLKQAAKSRRDNEVLKELVKKIRKVVNDAEDSIDKFVIEAKRHDDKNKFAQWFHLTHVARAKGVADEIKTIRERVKE
N4    MADVAVKFLVENLMQLLIDNADLILGIKGEVEDLLRDLNDFNAFLKQAAKSRRDNEVLKEMVKKIRKVVNDAEDSIDKFVIEAKRHDDKNKFAQWFHLTHVARAKGVADEIKSIRERVKE
N88   MADVAVKFLVENLMQLLIDNADLIIGIKGEVENLLQDLNDFNAFLKQAAKSRRDNEVLKSLVKKIRKVVNDAEDSIDKFVIEAKRHDDKNKFAQWFHLTHVARAKGVADEIKTIRERVKE
N95   MADVAVKFLVENLMQLLIDNAGLIIGIKGEVENLLQDLNDFNAFLKQAAKSRRDNEVLKSLVKKIRKVVNDAEDSIDKFVIEAKRHDDKNKFAQWFHLTHVARAKGVADEIKTIRERVKV
N89   MADVAVKFLVENLMQLLIDNADLIIGIKGEVENLLQDLNDFNAFLKQAAKSRRDNEVLKSLVKKIRKVVNDAEDSIDKFVIEAKRHDDKNKFAQWFHLTHVARAKGVADEIKTIRERVKE
      *********************.**:*******:**:***********************.:***************************************************:******

N3    IRQNDAYGLQAITFDDNSNRGAEERKVPVVEEDDVVGFDEEAKTVIDRLIGGSDDLEVVPVVGMPGLGKTTLAYKIYKDPKVEYEFFTRIWVYVSQSFKRREIFLNIISKFTRNTKQYHD
N4    IRENDSYGLQAIALDDNSNRGDEERKVPVVEEDDVVGFDEEAKTVIDRLIGGSDDVEVVPVVGMPGLGKTTLAYKIYKDPKVEYEFFTRIWVYVSQTFKRREIFLNIISKFTRNTKQYHD
N88   IRQNDAYGLQAITSDDNSNKGAQERKVPVVEEDDVVGFDEEAKTVIDRLIGGSDYLEVVPVVGMPGLGKTTLAYKIYKDPKVEYEFFTRIWVYVSQSFKRREIFLNIISKFTRNTKQYHD
N95   IRQNDAYGLQAITSDDNSNKGAQERKVPVVEEDDVVGFDEEAKTVIDRLIGGSDHLEVVPIVGMPGLGKTTLAYKIYKDPKVECEFFPCIWVNVSQLYKRREIFLNIISKFTRNTKQYYD
N89   IRQNDAYGLQAITSDDNSNQGAQERKVPVVEEDDVVGFDEEAKTVIDRLIGGSDY--VVPVVGMPGLGKTTLAYKIYKDPKVEYEFFNRIWVYVSQSFNRREIFLNIISKFTRNTKQYHD
      **:**:******: *****:* :*******************************   ***:********************** ***  *** *** ::*******************:*

N3    TPEEDLANEVKELLGKGGKYLIVLDDVWTMEAWDRIKIAFPNNGKRNRVLMTTRQSNVAKCCNDKPHDLKFLTENESWELLEKKVFHKEKCPPELELPGKSIAEKCRGLPLAIVVIAGAL
N4    TPEEDLANEVKELLGKGGKYLIVLDDVWTMEAWDRIKIAFPNNGKRNRVLMTTRQTNVAKCCNDKPHDLKFLTKDESWELLEKKVFHKEKCPPELELPGKSIAEKCRGLPLAIVVIAGAL
N88   TPEEDLANEVKELLGKGGKYLIVLDDVWTMEAWDRIKIAFPNNGKRNRVLMTTRQSNVAKCCNDKPHDLKFLTENESWELLEKRVFHKEKCPPELELPGKSIAEKCRGLPLAIVVIAGAL
N95   VPEEDLADEARELLGKGGKYLIVLDDVWTMEAWDRIKIAFPNNGKCSRVLMTTRQSNVARYCNDKPHDLKFLTENESWELLEKRVFHKEKCPPELELPGKSIAEKCRGLPLAIVVIAGAL
N89   TPEEELANEIKELLGKGGKYLIVLDDVWTREAWDRIKIAFPNNDKRNRVLMTTRQSNVAKCCNDKPHDLKFLTENESWELLEKRVFHKEKCPPELELPGKSIAEKCRGLPLAIVVIAGAL
      .***:**:* :****************** *************.* .********:***: ************:.********:************************************

N3    IGKGKTTREWELVADSVGEHLINRDPENCKKLVQMSYDRLPYDLKACFLYCGAFPGGSEIPARKLIRLWIAEGFIQYQGPLTLEDIAEDHLNDLVNRNLVMVMQRSSSGQIKTCRVHDML
N4    IGKGKTTREWELVADSVGEHLINRDPENCKKLVQMSYDRLPYDLKACFLYCGAFPGGSQIPARKLIRLWIAEGFIQYQGPLTLEDVAEDHLNDLVNRNLVMVMQRSCSGQIKTCRVHDML
N88   IGKGKTTREWELVADSVGEHLINRDPENCKKLVQMSYDRLPYDLKACFLYCGAFPGGSEISARKLICLWIAEGFIQYQGPLTLEDIAEDHLNDLVNRNLVMVMQRSSSGQIKTCRVHDML
N95   IGKGKTTREWELVADSVGEHLINRDPENCKKLVQMSYDRLPYDLKACFLYCGAFPGGSEISARKLICLWIAEGFIQYQGPLTLEDIAEDHLNDLVNRNLVMVMQRSSSGQIKTCRVHDML
N89   IGKGKTTREWELVADSVGEHLINRDPENCKKLVQMSYDRLPYDLKACFLYCGAFPGGSEISARKLICLWIAEGFIQYQGPLTLEDIAEDHLNDLVNRNLVMVMQRSSSGQIKTCRVHDML
      **********************************************************:*.***** ******************:********************.*************

N3    HEFCRHEAMMEENLFQEIKRGQEQSFPEKQELATYRRLCIHSSVSEFLSTKPFGEHVRSFLCFASKKIEMPSGEIPAIPKAFPLLRVLDAESIKFSRFSKEFFKLFHLRYIAFSTDSIKT
N4    HEFCRHEAMMEENLFQEIKRGQERSFPEKQELATYRRLCIHSSVPEFLSTKPSGEHVRSFLCFGSKKIDMPSNEIPVIPKAFPLLRVLDAESIKFNRFSREFFKLFHLRYIAFSTDKIKT
N88   HEFCRHEAMMEENLFQEIKRGQEHSFPEKQELATYRRLCIHSSVSEFLSTKPFGEHVRSFLCFASKKFEMPLGEIPAIPRAFPLLRVLDAESIKFSRFSKEFFKLFHLRYIAFSTDSIMT
N95   HEFCRHEAMMEENLFQEIKRGQEHSFPEKQELATYRRLCIHSSVSEFLSTKPFGEHVRSFLCFASKKFEMPLGDI--IPRAFPLLRVLDAESIKFSRFPKEFFKLFHLRYIAFSTDSIMT
N89   HEFCRHEAMMEENLFQEIKRGQEHSFPEKQELATYRRLCIHSSVSEFLSTKPFGEHVRSFLCFASKKFEMPLGEIPAIPRAFPLLRVLDAESIKFSRFSKEFFKLFHLRYIAFSTDSIMT
      ***********************:********************.******* **********.***::** .:*  **:***************.**.:****************.* *

N3    IPTNIGNLWNVQTLIIETQQGTLDIKADIWNMTRLRHVCTNASATLPSPKRPKSSKDNLVNRCLQTLSTIAPECCTAEVFTRTPNLKKLGVRGKIAALLETSKDGSGSGLFSNIGKLDCL
N4    IPTDIGNLWNVQTLIVETQQGTLDIKADIWNMTRLRHVCTNASATLPSTKRPKSSKDNLANRCLQTLSTIAPECCTAEVFTRTPNLKKLGVRGKIAALLETSKDGSGSGLFSNIGKLACL
N88   IPTNIGNLWNVQTLIIETQQGTLDIKADIWNMTRLRHVCTNASATLPSPKRPKSSKDNLVNRCLQTLSTIAPECCTAEVFTRTPNLKKLGVRGKIAALLETSKDGSGSGLFSNIGKLDCL
N95   IPTNIGNLWNVQTLIIETQQGTLDIKADIWNMTRLRHVCTNASAILPSPKHPKSRKE--VNHYLQTLSTIAPECCTAEIFTRTPNLKKLGVRGKIAALLETSKDGSGSGLFSNIGKLDCL
N89   IPTNIGNLWNVQTLIIETQQGTLDIKADIWNMTRLRHVCTNASATLPSPKRPKSSKDNLVNRCLQTLSTIAPECCTAEVFTRTPNLKKLGVRGKIDALLETSKDGSSSGLFSNIGKLDCL
      ***:***********:**************************** ***.*:*** *:  .*: ***************:**************** **********.********** **

N3    ENLKLVNDTRLSSKQLHLPPAYIFPQKLKKLTLIDTWFEWKDMSILGLLEYLEVLKLKENAFRGQSWEPEDGGFPRLQVLWIERTDLSSWKASSGNFPRLKRLVLISCDKLEELPAELAD
N4    ENLKLVNDTRLSSKQLHLPPAYIFPQKLKKLTLVDTWFEWKDMSILGLLEYLEVLKLKENAFRGQSWEQEDGGFPRLQVLWIERTDLTSWKASSGNFPRLKHLVLISCDKLEELPAELAD
N88   ENLKLVNDTRQSSKQLHLPPAYIFPQKLKKLTLIDTWFEWKDMSILGLLEYLEVLKLKENAFRGQSWEPEDGGFPRLQVLWIERTDLSSWKASSGNFPRLKRLVLISCDKLEELPAELAD
N95   ENLKLVNDTRQSGKQLHLPPAYIFPQKLKKLTLIDTWFEWKDMSILGLLEYLEVLKLKENAFRGQSWEPEDGGFPRLQVLWIERTDLSSWKASSGNFPQLKRLVLISCDKLEELPAALAD
N89   EKLKLVNDTRQSRKQLHLPPAYIFPQKLKKLTLIDTWFEWKDMSILGLLEYLEVLKLKENAFRGQSWEPEDGGFPRLQVLWIERTDLSSWKASSGNFPRLKRLVLIACDNLKELPAELAD
      *:******** * ********************:********************************** ******************:**********:**:****:**:*:**** ***

N3    VQNLQLMELQSTSESAAKSARAILKKKQEKEQDGTKSSGFKLSIFPPDLGL
N4    VKNLQLMELQSTSESAARSARAILKKKQEKEQDGSKSSGFKLSIFPPDLGL
N88   VQNLQLMELQSTSESAAKSARAILKKKQEKEEDGTKSSGFKLSVFPPDLGL
N95   VQSLQLMELHYTSESAAKSARAILKKRLRKEEGSTNRCGCKLSVFPPDLGL
N89   VENLQLMELQSTSVSAAKSARAILKKKQEKE-----GSGFKLSVFPPDLGL
      *:.******: ** ***:********: .**      .* ***:*******
```

**Fig. S5.**

Amino acid sequence alignment of ancestral NRC3 variants. The alignment was done using MAFFT (Multiple Alignment using Fast Fourier Transform). N3: the ancestral variant before the divergent of three allelic groups. N4: the ancestral variant of NRC3a. N88: the ancestral variant before divergent of NRC3b/c. N95: the ancestral variant of NRC3b. N89: the ancestral variant of NRC3c.

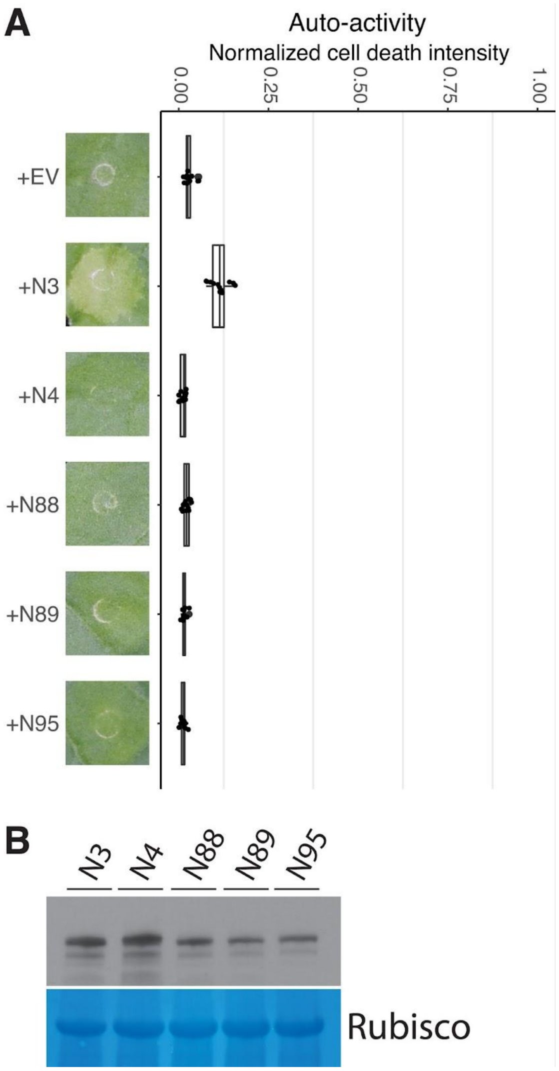


**Fig. S6.**

NRC3 ancestral variants show no or low auto-activities when expressed in *N. benthamiana*. (A) Auto-activity analysis of ancestral NRC3 variants. The NRC3 variants were expressed alone in WT *N. benthamiana.* Cell death intensity and phenotypes were recorded at 6 dpi. The dot plots represent cell death intensity quantified using UVP ChemStudio PLUS. (B) Protein accumulation of ancestral NRC3 variants. NRC3 variants were transiently expressed in WT *N. benthamiana.* The proteins were extracted from leaf samples at 2 dpi and the NRC3 protein accumulations were detected by α-myc antibody. SimplyBlue SafeStain-staining of Rubisco was used as the loading control.

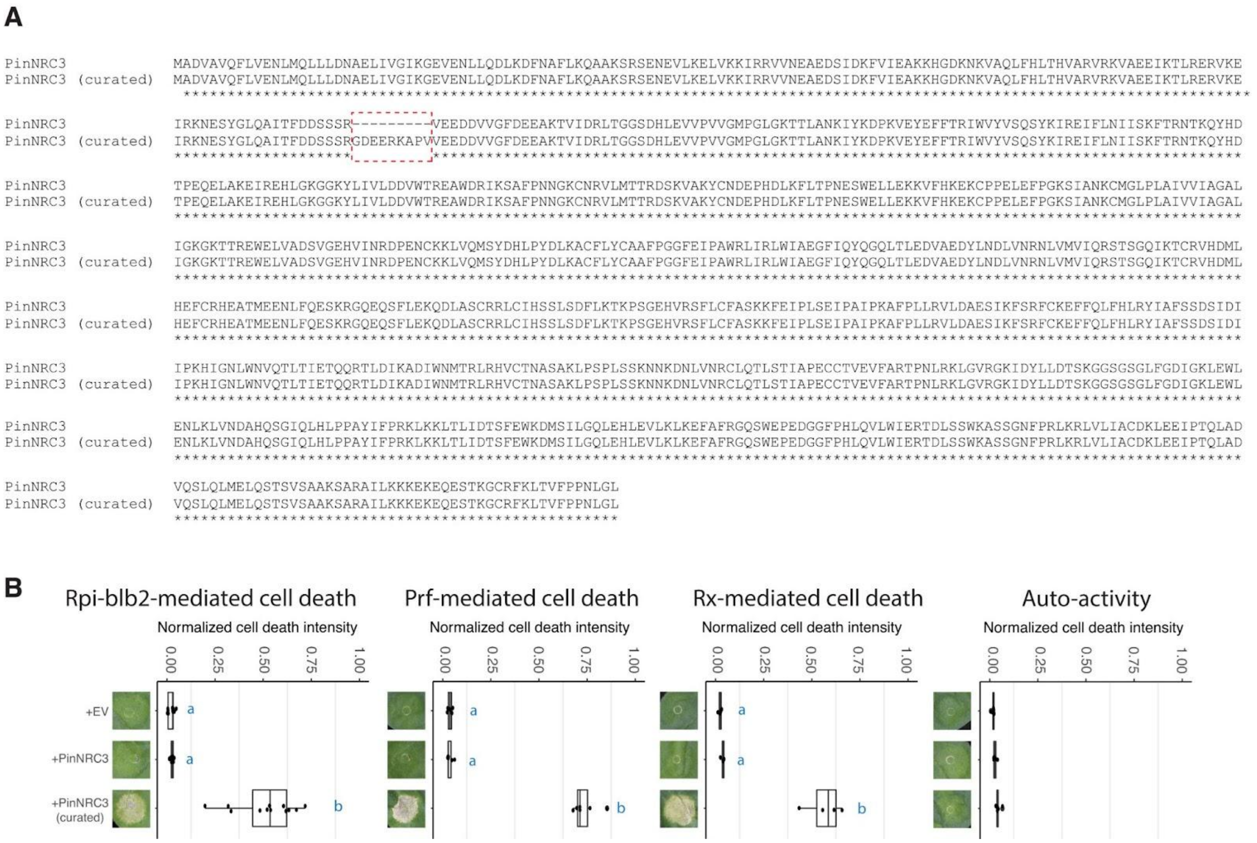


**Fig. S7.**

The manually curated PinNRC3 variant rescued Rpi-blb2/Prf/Rx-mediated cell death. (A) Amino acid sequence comparison of *Petunia* NRC3 from the genome database and curated version. The indel was curated using sequences from SlNRC3. (B) Cell death assay of PinNRC3 variants co-expressed with Rpi-blb2/AVRblb2, Pto/AvrPto, or Rx/CP in *nrc2/3/4*_KO *N. benthamiana*. The auto-activity analysis was done by expressing NRC3 variants alone in WT *N. benthamiana*. The dot plots represent cell death intensity quantified by UVP ChemStudio PLUS at 6dpi. Statistical differences were analyzed with Tukey's HSD test ($p<0.05$).

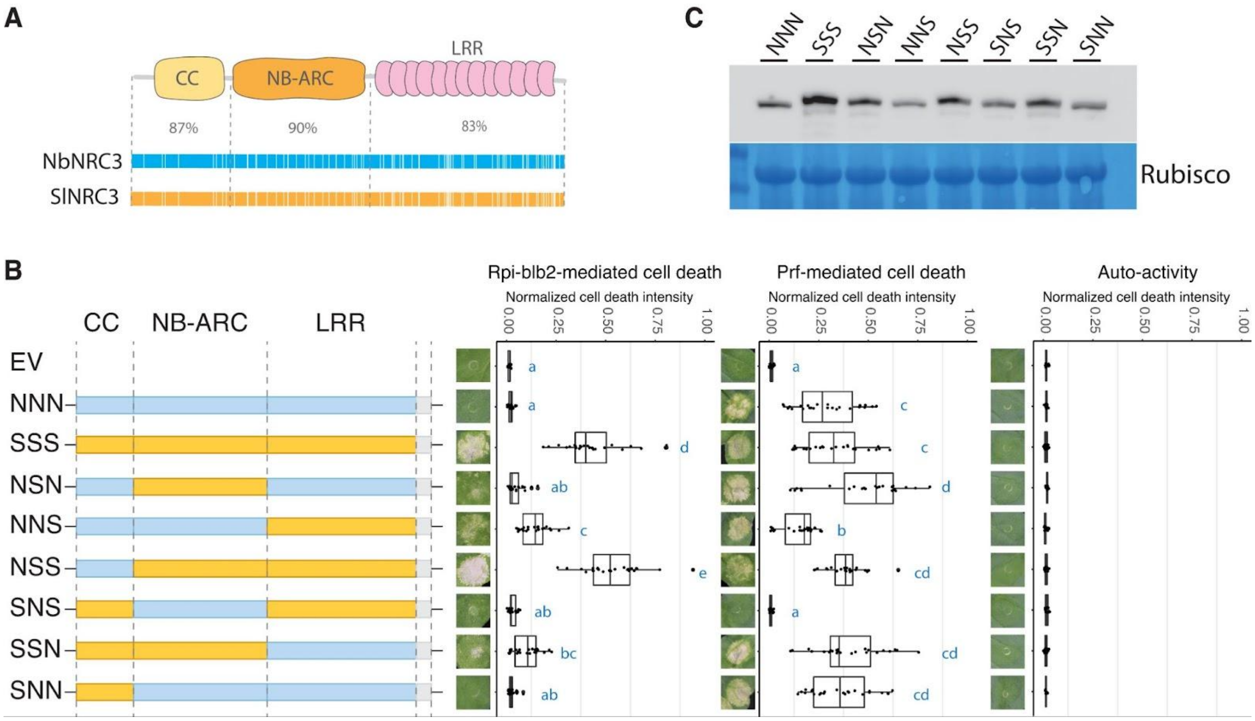


**Fig. S8.**

The NB-ARC and LRR domains cooperatively contributed to the helper-sensor compatibility. (A) Comparison of three major domains between NbNRC3 and SlNRC3. SlNRC3 was painted in orange while NbNRC3 was painted in blue. The two NRC variants shared around 86.42% overall sequence identity, with 87%, 90%, and 83% for the CC domain, NB-ARC domain, and LRR domain, respectively. (B) Cell death assay of chimeric NbNRC3/SlNRC3 variants. The NRC3 chimeric variants were co-expressed with Rpi-blb2/AVRblb2 or Pto/AvrPto in *nrc2/3/4*_KO *N. benthamiana*. Auto-activity analysis was done by expressing NRC3 variants alone in WT *N. benthamiana*. The dot plots represent cell death intensity quantified by UVP ChemStudio PLUS at 6 dpi. Statistical differences were analyzed with Tukey's HSD test ($p<0.05$). (C) Protein accumulation of chimeric NRC3 variants tested in (B). NRC3 variants were transiently expressed in WT *N. benthamiana.* The proteins were extracted from leaf samples at 2 dpi and the NRC3 protein accumulations were detected by α-myc antibody. SimplyBlue SafeStain-staining of Rubisco was used as the loading control.

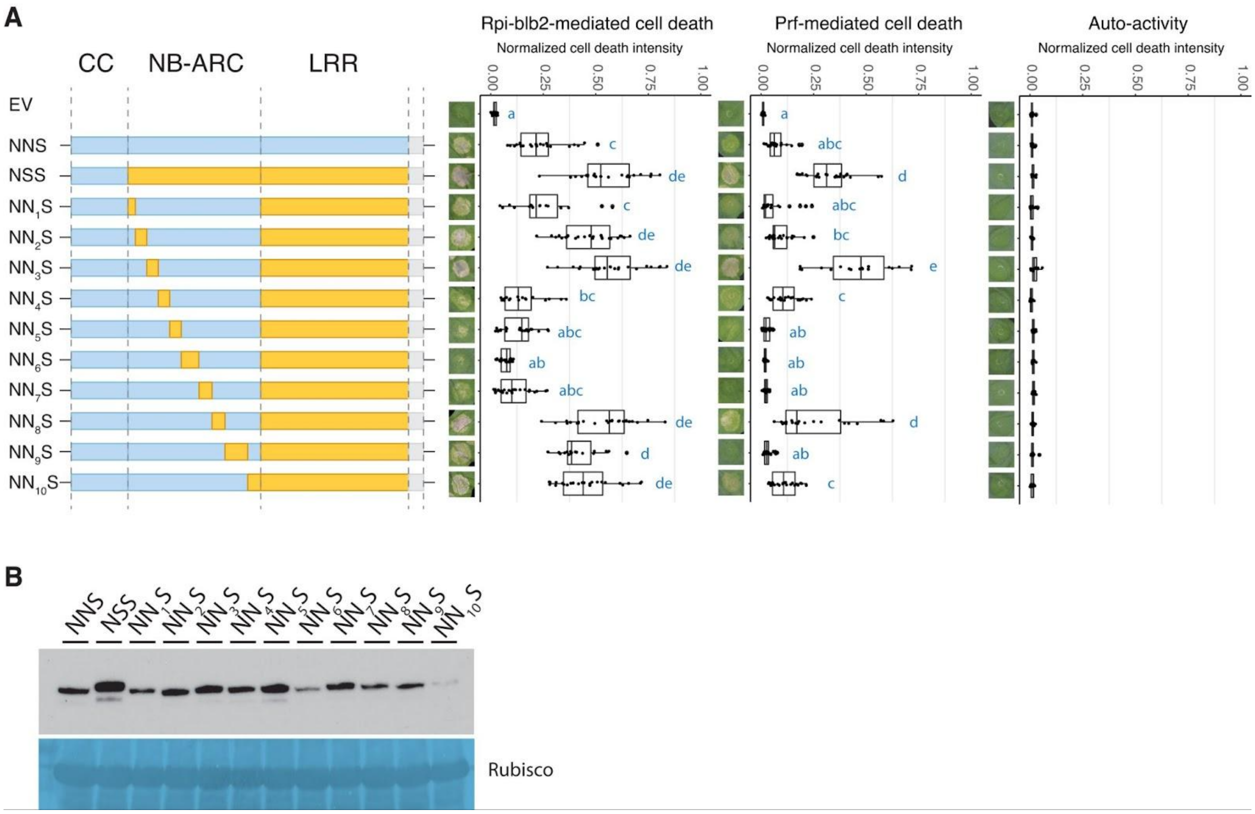


**Fig. S9.**

Region 3 of the NB-ARC domain contributed to the helper-sensor compatibility. (A) Cell death assays of chimeric NRC3 variants designed for investigating regions of the NB-ARC domain that contribute to helper-sensor compatibility. The variants were co-expressed with Rpi-blb2/AVRblb2 or Pto/AvrPto in *nrc2/3/4*_KO *N. benthamiana*. Auto-activity analysis was done by expressing NRC3 variants alone in WT *N. benthamiana*. The dot plots represent cell death intensity quantified by UVP ChemStudio PLUS at 6 dpi. Statistical differences were analyzed with Tukey's HSD test ($p<0.05$). (B) Protein accumulation of chimeric NRC3 variants tested in (A). NRC3 variants were transiently expressed in WT *N. benthamiana.* The proteins were extracted from leaf samples at 2 dpi and the NRC3 protein accumulations were detected by α-myc antibody. SimplyBlue SafeStain-staining of Rubisco was used as the loading control.

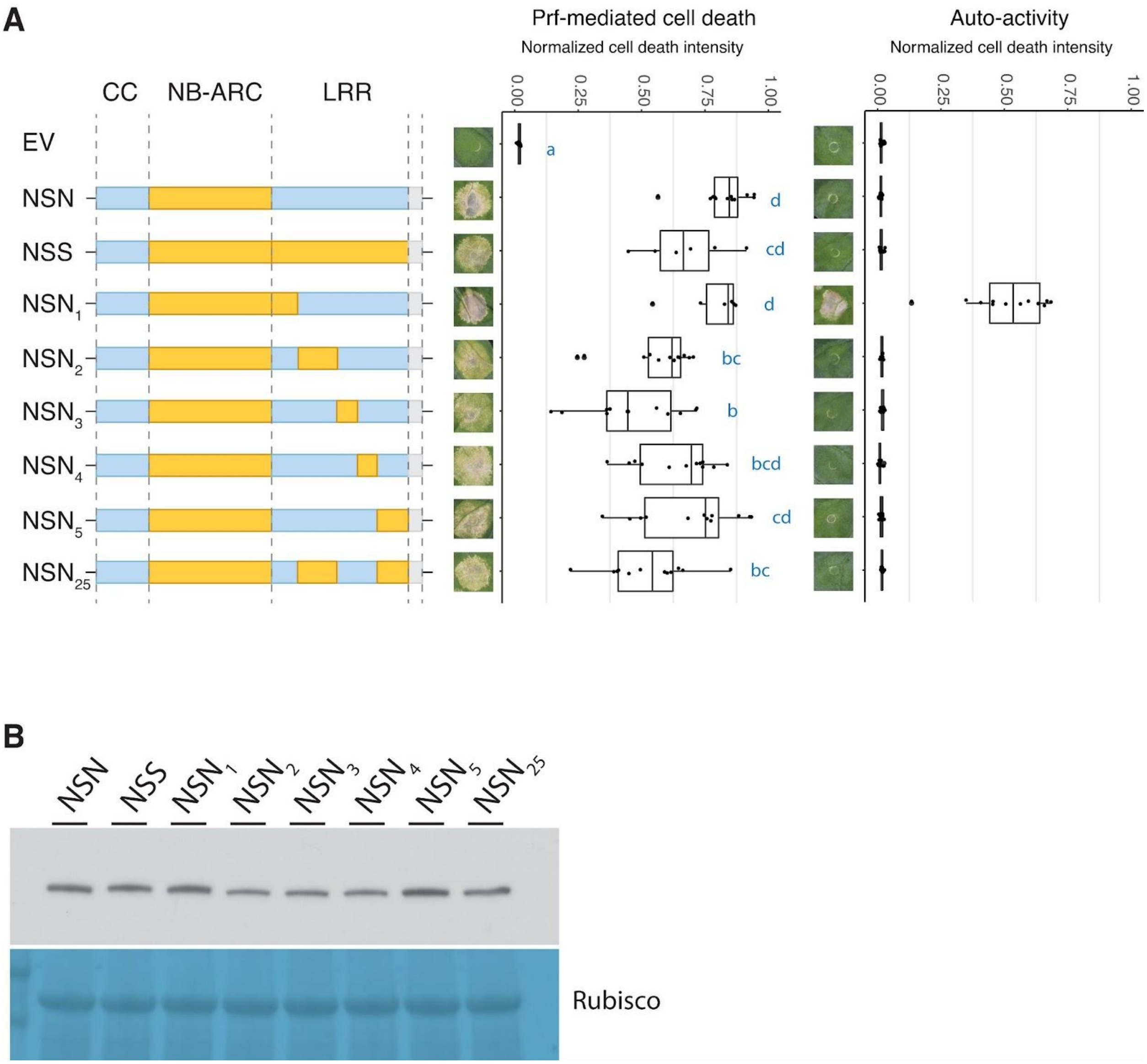


**Fig. S10.**

NRC3 variants $NSN_2$ to $NSN_5$ and $NSN_{25}$ functioned with Prf. (A) Cell death assays of chimeric NRC3 variants tested in Fig. 3B. The variants were co-expressed with Pto/AvrPto in *nrc2/3/4*_KO *N. benthamiana*. Auto-activity analysis was done by expressing NRC3 variants alone in WT *N. benthamiana*. The dot plots represent cell death intensity quantified by UVP ChemStudio PLUS at 6dpi. Statistical differences were analyzed with Tukey's HSD test ($p<0.05$). (B) Protein accumulation of chimeric NRC3 variants tested in (A). NRC3 variants were transiently expressed in WT *N. benthamiana.* The proteins were extracted from leaf samples at 2 dpi and the NRC3 protein accumulations were detected by α-myc antibody. SimplyBlue SafeStain-staining of Rubisco was used as the loading control.

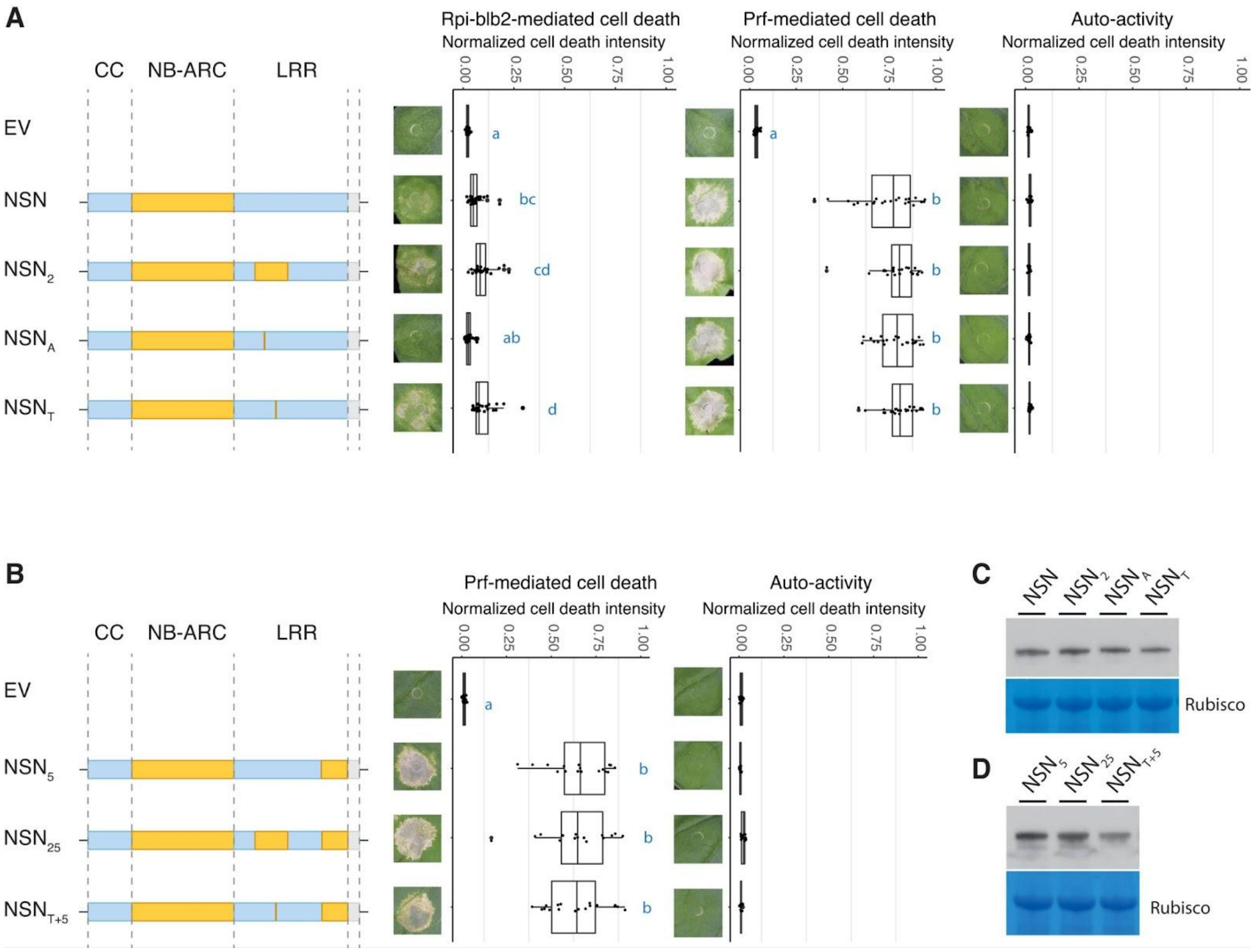


### Fig. S11.

A single amino acid change (I642T) conferred full activity in rescuing Rpi-blb2 cell death in the $NSN_5$ background. (A) Cell death assay of chimeric NRC3 variants carrying T607A or I642T in NSN background. The variants were co-expressed with Rpi-blb2/AVRblb2 or Pto/AvrPto in *nrc2/3/4*_KO *N. benthamiana*. (B) Cell death assay of chimeric NRC3 variants tested in Fig. 3C. The variants were co-expressed with Pto/AvrPto in *nrc2/3/4*_KO *N. benthamiana*. Auto-activity analysis was done by expressing NRC3 variants alone in WT *N. benthamiana*. The dot plots represent cell death intensity quantified by UVP ChemStudio PLUS at 6 dpi. Statistical differences were analyzed with Tukey's HSD test ($p<0.05$). (C and D) Protein accumulation of chimeric NRC3 variants tested in (A) and (B). NRC3 variants were transiently expressed in WT *N. benthamiana.* The proteins were extracted from leaf samples at 2 dpi and the NRC3 protein accumulations were detected by α-myc antibody. SimplyBlue SafeStain-staining of Rubisco was used as the loading control.

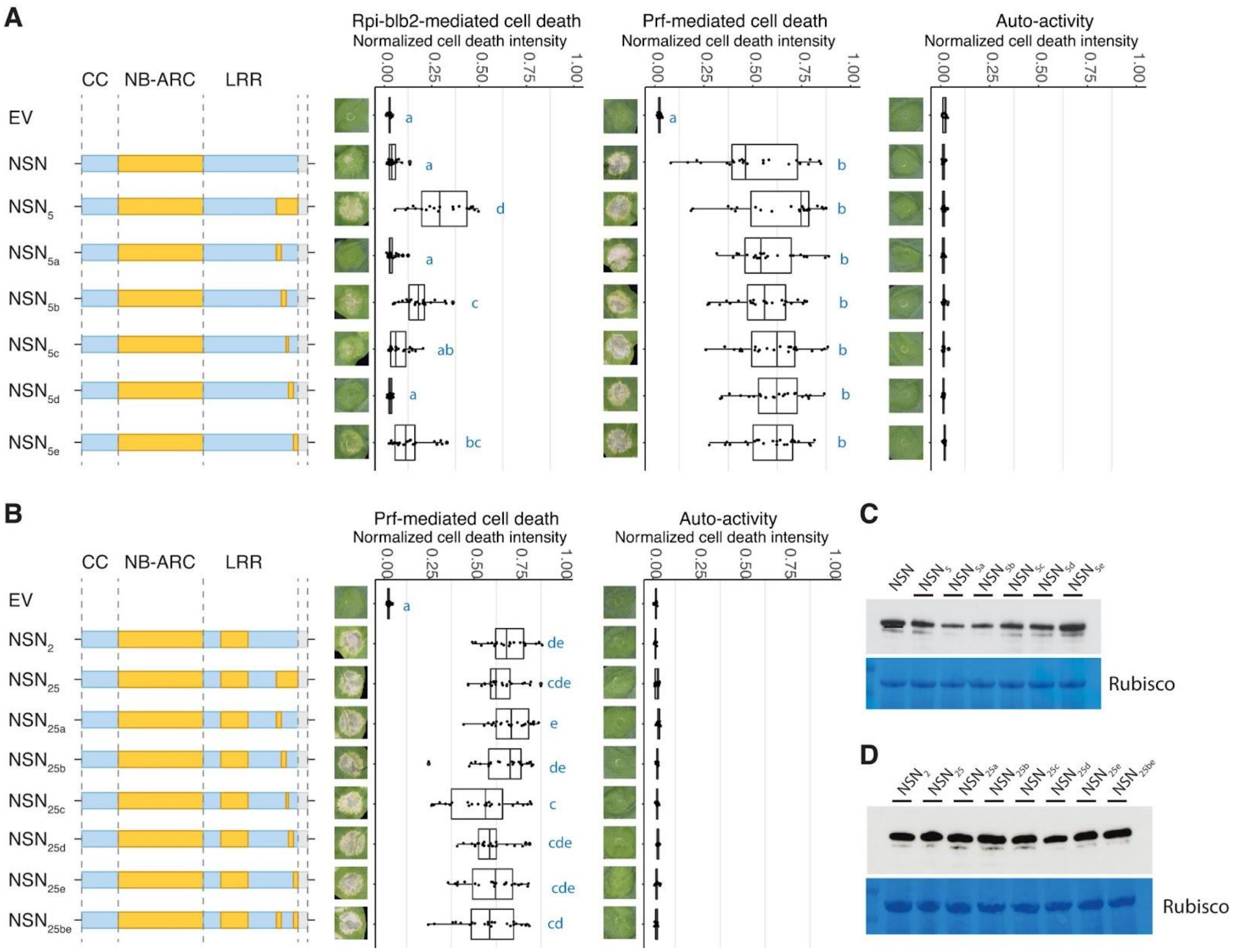


**Fig. S12.**

The NRC3 variant $NSN_{25be}$ fully rescued the Rpi-blb2-mediated cell death in the *nrc2/3/4*_KO *N. benthamiana* (A) Cell death assay of chimeric NRC3 variants carrying LRR region 5a to 5e of SlNRC3 in NSN background. The variants were co-expressed with Rpi-blb2/AVRblb2 or Pto/AvrPto in *nrc2/3/4*_KO *N. benthamiana*. (B) Cell death assay of chimeric NRC3 variants tested in Fig. 3D. The variants were co-expressed with Pto/AvrPto in *nrc2/3/4*_KO *N. benthamiana*. Auto-activity analysis was done by expressing NRC3 variants alone in WT *N. benthamiana*. The dot plots represent cell death intensity quantified by UVP ChemStudio PLUS at 6 dpi. Statistical differences were analyzed with Tukey's HSD test ($p<0.05$). (C) and (D) Protein accumulation of chimeric NRC3 variants tested in (A) and (B). NRC3 variants were transiently expressed in WT *N. benthamiana.* The proteins were extracted from leaf samples at 2 dpi and the NRC3 protein accumulations were detected by α-myc antibody. SimplyBlue SafeStain-staining of Rubisco was used as the loading control.

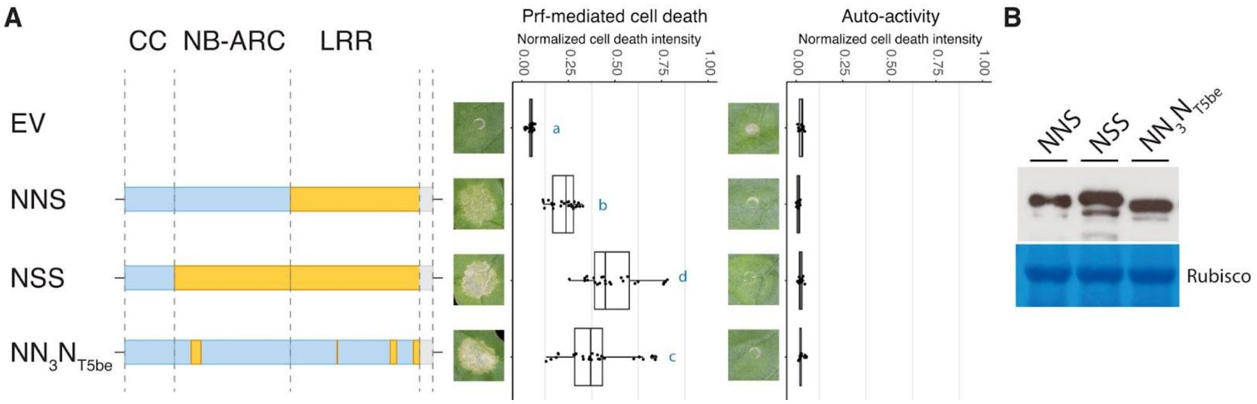


**Fig. S13.**

$NN_3N_{T5be}$ functioned with Prf and was not auto-active. (A) Cell death assay of chimeric NRC3 variants tested in Fig. 3E. The variants were co-expressed with Pto/AvrPto in *nrc2/3/4*_KO *N. benthamiana*. Auto-activity analysis was done by expressing NRC3 variants alone in WT *N. benthamiana*. The dot plots represent cell death intensity quantified by UVP ChemStudio PLUS at 6dpi. Statistical differences were analyzed with Tukey's HSD test ($p<0.05$). (B) Protein accumulation of chimeric NRC3 variants tested in (A). NRC3 variants were transiently expressed in WT *N. benthamiana.* The proteins were extracted from leaf samples at 2 dpi and the NRC3 protein accumulations were detected by α-myc antibody. SimplyBlue SafeStain-staining of Rubisco was used as the loading control.

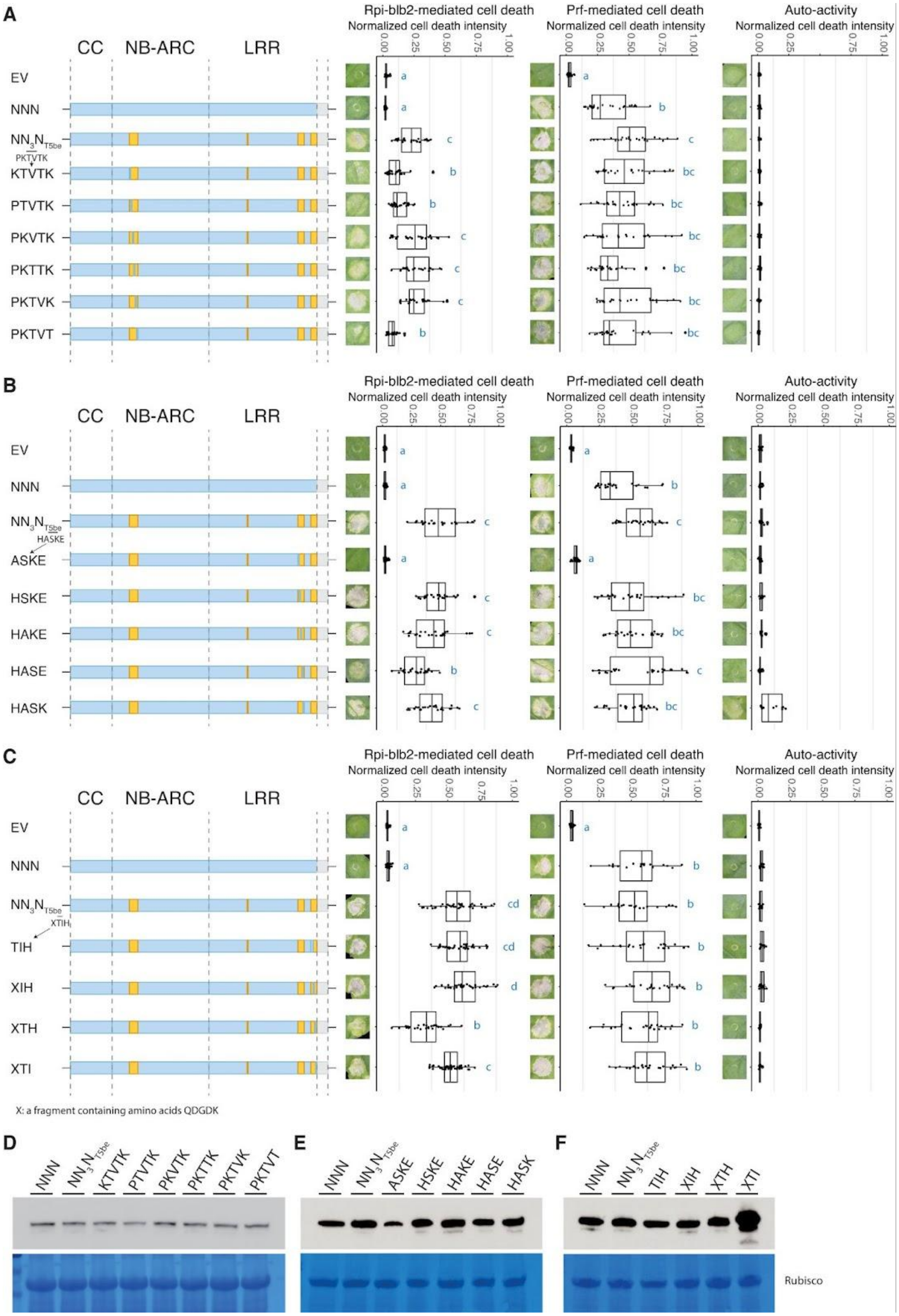
A
B
C
D
E
F
Rpi-blb2-mediated cell death
Prf-mediated cell death
Auto-activity
Normalized cell death intensity
CC
NB-ARC
LRR
EV
NNN
PKTVTK
KTVTK
PTVTK
PKVTK
PKTTK
PKTVK
PKTVT
HASKE
ASKE
HSKE
HAKE
HASE
HASK
XTIH
TIH
XIH
XTH
XTI
X: a fragment containing amino acids QDGDK
Rubisco

**Fig. S14.**

NRC3 variants with S202P, T203K, N221K, C824H, N832K, or V881I mutations quantitatively affected the ability of NRC3 to function with Rpi-blb2. Cell death assay of chimeric NRC3 variants designed for pinpointing the polymorphisms in (A) NB-ARC domain region 3 which contains six amino acid differences, (B) LRR domain region 5b containing five amino acid differences, and (C) LRR domain region 5e contain a small insertion/indel (labeled as X) three amino acid (TIH) differences between NbNRC3c and SlNRC3a. The analysis was done in the $NN_3N_{T5be}$ background by replacing each residue with the amino acid of NbNRC3c. The variants were co-expressed with Rpi-blb2/AVRblb2 or Pto/AvrPto in *nrc2/3/4*_KO *N. benthamiana*. Mutations at positions 202, 203, 221 (NB-ARC domain), 824, 832 (LRR domain region 5b), and 881 (LRR domain region 5e) affected the ability of NN3NT5be to function with Rpi-blb2. Auto-activity analysis was done by expressing NRC3 variants alone in WT *N. benthamiana*. The dot plots represent cell death intensity quantified by UVP ChemStudio PLUS at 6 dpi. Statistical differences were analyzed with Tukey's HSD test ($p<0.05$). (D to F) Protein accumulation of chimeric NRC3 variants tested in (A to C). The proteins were extracted from leaf samples at 2 dpi and the NRC3 protein accumulations were detected by α-myc antibody. SimplyBlue SafeStain-staining of Rubisco was used as the loading control.

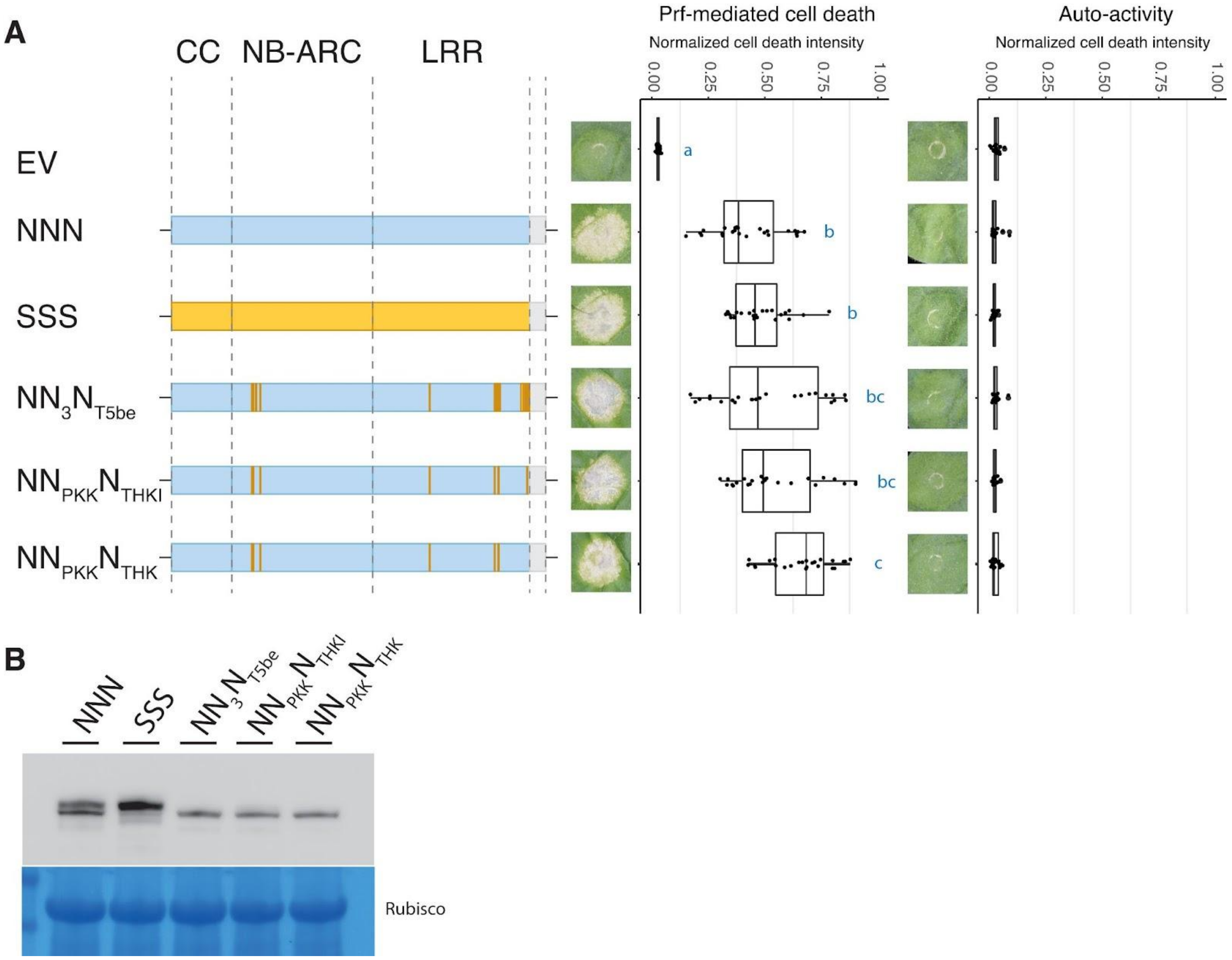


**Fig. S15.**

$NN_{PKK}N_{THKI}$ and $NN_{PKK}N_{THK}$ functioned with Prf and were not auto-active. (A) Cell death assay of chimeric NRC3 variants tested in Fig. 3G. The variants were co-expressed with Pto/AvrPto in *nrc2/3/4*_KO *N. benthamiana*. Auto-activity analysis was done by expressing NRC3 variants alone in WT *N. benthamiana*. The dot plots represent cell death intensity quantified by UVP ChemStudio PLUS at 6 dpi. Statistical differences were analyzed with Tukey's HSD test ($p<0.05$). (B) Protein accumulation of chimeric NRC3 variants tested in (A). The proteins were extracted from leaf samples at 2 dpi and the NRC3 protein accumulations were detected by α-myc antibody. SimplyBlue SafeStain-staining of Rubisco was used as the loading control.

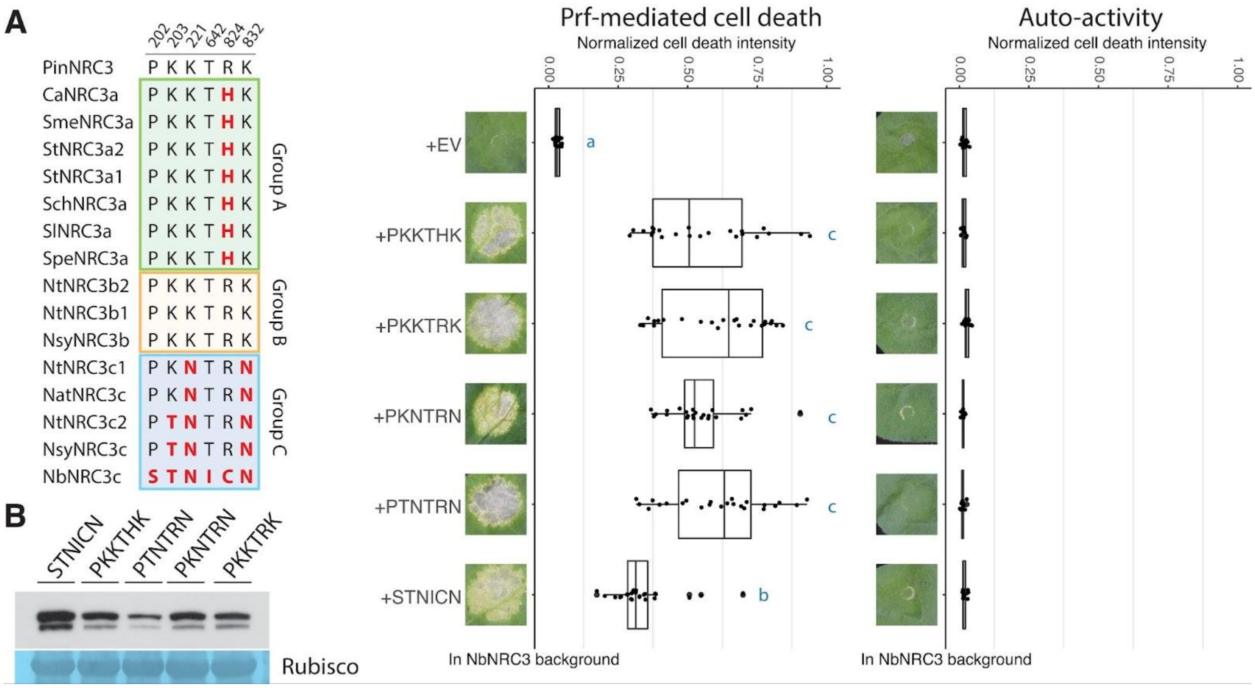


**Fig. S16.**

The NbNRC3 variants carrying polymorphisms from three allelic groups functioned with Prf and were not auto-active. (A) Cell death assays of NRC3 variants tested in Fig. 4A. Left panel, the polymorphisms of NRC3 natural variants at the six positions. The middle and right panels, the cell death assays and the auto-activity analysis. The variants were co-expressed with Pto/AvrPto in *nrc2/3/4*_KO *N. benthamiana*. Auto-activity analysis was done by expressing NRC3 variants alone in WT *N. benthamiana*. The dot plots represent cell death intensity quantified by UVP ChemStudio PLUS at 6 dpi. Statistical differences were analyzed with Tukey's HSD test ($p<0.05$). (B) Protein accumulation of NRC3 variants tested in (A). The proteins were extracted from leaf samples at 2 dpi and the NRC3 protein accumulations were detected by α-myc antibody. SimplyBlue SafeStain-staining of Rubisco was used as the loading control.

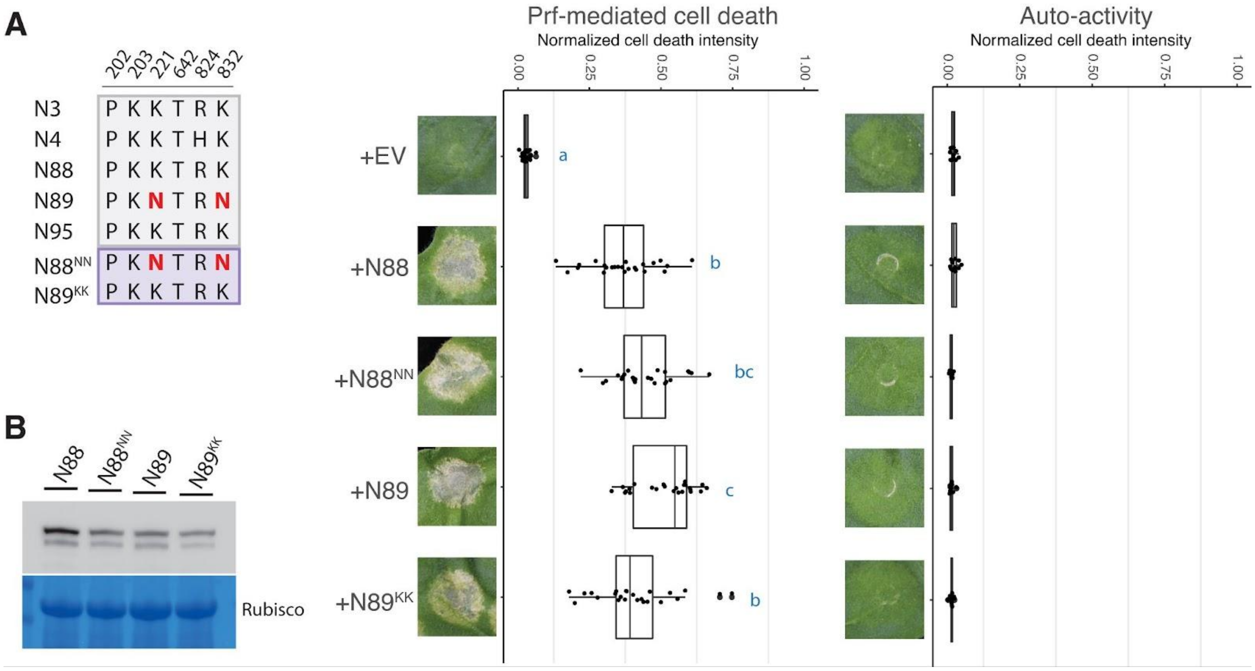


**Fig. S17.**

NRC3 variants N88NN and N89KK functioned with Prf and were not auto-active. (A) Cell death assay of NRC3 variants tested in Fig. 4B. Left panel, the polymorphisms of ancestral NRC3 variants at the six positions. The middle and right panels, the cell death assays and the auto-activity analysis. The variants were co-expressed with Pto/AvrPto in *nrc2/3/4*_KO *N. benthamiana*. Auto-activity analysis was done by expressing NRC3 variants alone in WT *N. benthamiana*. The dot plots represent cell death intensity quantified by UVP ChemStudio PLUS at 6 dpi. Statistical differences were analyzed with Tukey's HSD test ($p<0.05$). (B) Protein accumulation of NRC3 variants tested in (A). The proteins were extracted from leaf samples at 2 dpi and the NRC3 protein accumulations were detected by α-myc antibody. SimplyBlue SafeStain-staining of Rubisco was used as the loading control.

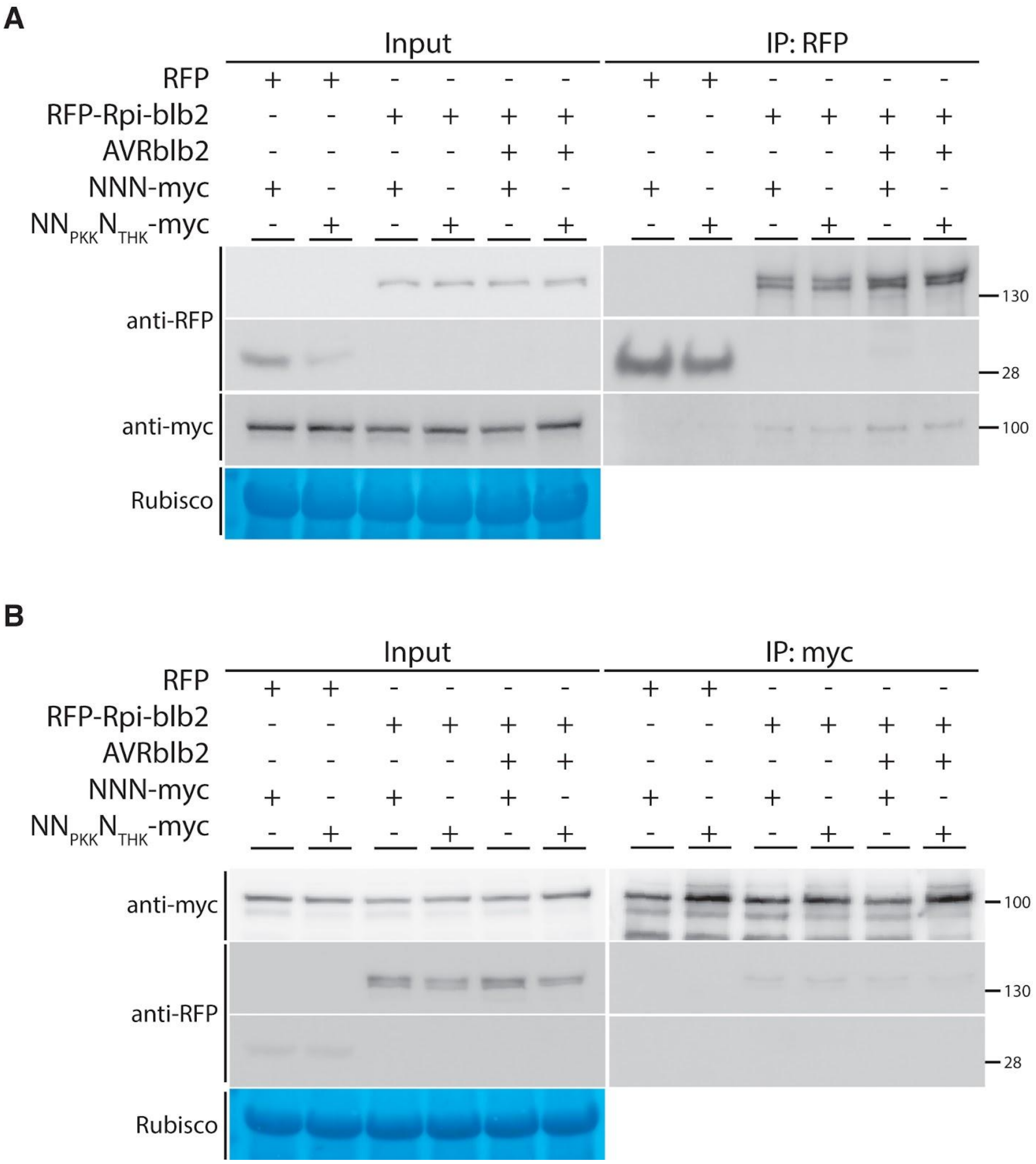


**Fig. S18.**

Steady-state interactions detected using co-IP did not reflect the compatibility between Rpi-blb2 and NRC3 variants. Co-immunoprecipitation assay of Rpi-blb2 and NRC3 variants. (A) RFP-Rpi-blb2 and NRC3s-myc were co-expressed with or without flag-AVRblb2. NRC3s-myc coexpressed with RFP were used as negative controls. Protein extracts (input) and RFP-Trap pull-down samples (IP) were analyzed using Western blot analysis with α-RFP, and α-myc antibodies. SimplyBlue

SafeStain-staining of Rubisco was used as the loading control. (B) Protein extracts (input) and myc-Trap pull-down samples (IP) were analyzed using Western blot analysis using α-RFP, and α-myc antibodies. SimplyBlue SafeStain-staining of Rubisco was used as the loading control.

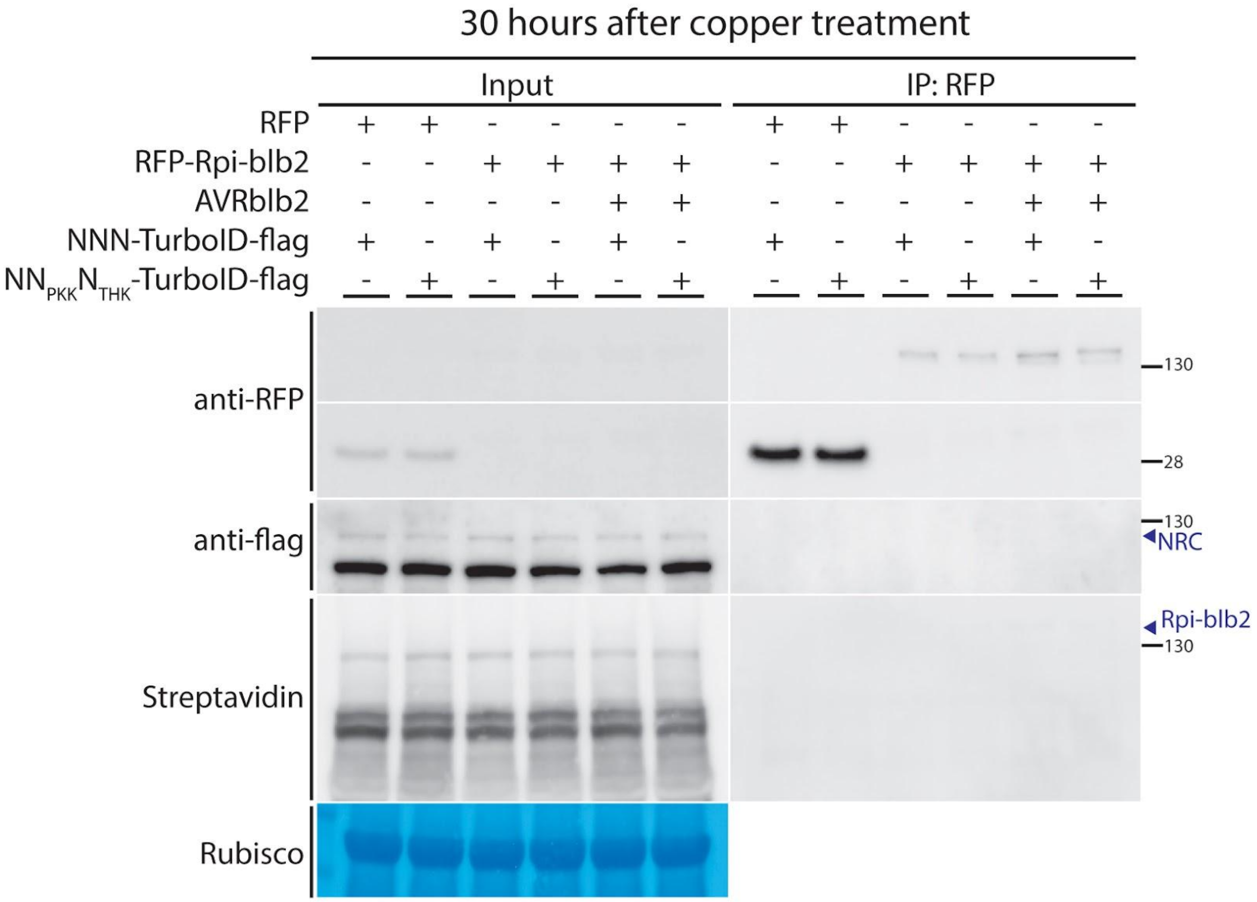


**Fig. S19.**

The level of biotinylation on Rpi-blb2 was barely detectable at 30 hours post-copper treatment. Western blot of TurboID-based proximity labeling assay of sample collected at 30 hours post-copper treatment. RFP-Rpi-blb2 and NRC3s-TurboID-flag were co-expressed in *nrc2/3/4*_KO *N. benthamiana*. NRC3s-myc coexpressed with RFP were used as negative controls. All treatments were coexpressed with copper inducible flag-AVRblb2 and CUP2-P65. Treatments with AVRblb2 induction were indicated with '+' while treatments without AVRblb2 induction were indicated with '-'. Protein extracts (input) and RFP-Trap pull-down samples (IP) were analyzed using Western blot analysis using α-RFP, α-flag antibodies, and streptavidine fused with HRP. SimplyBlue SafeStain-staining of Rubisco was used as the loading control.

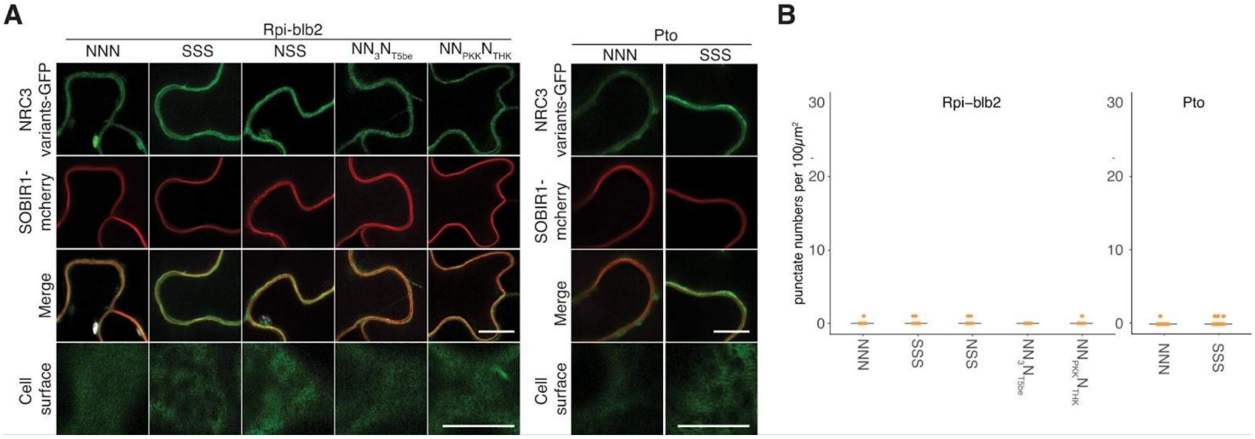


**Fig. S20.**

NRC3 variants do not form membrane-associated punctate dots in the presence of sensor NLRs without corresponding effectors. (A) NRCs-GFP were co-expressed with HF-Rpi-blb2 or Pto. Samples were examined at 3 dpi. Scale bars represent 10µm. SOBIR1-mcherry was used as a plasma membrane marker. (B) Quantification of punctate dots of NRCs in (A).

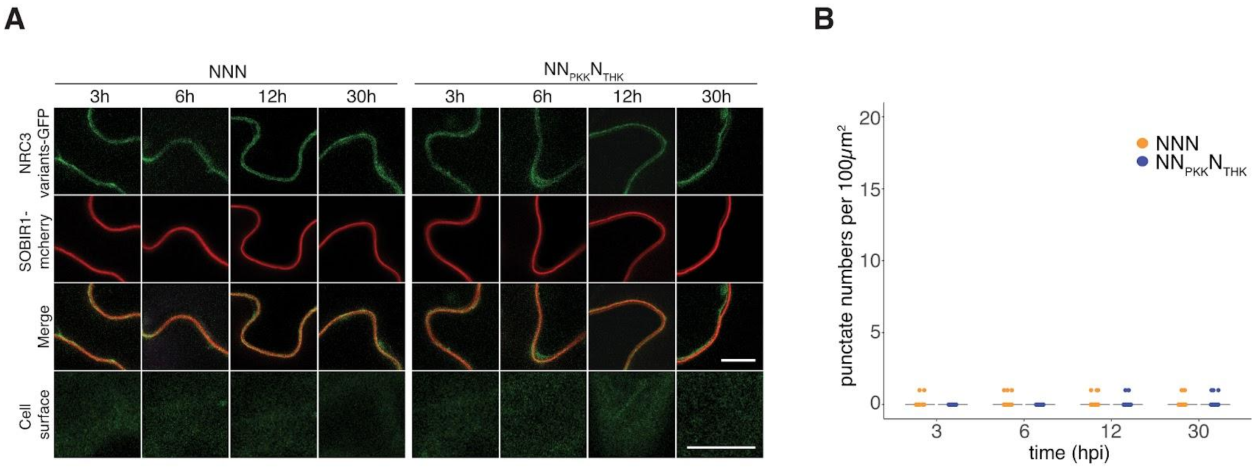


**Fig. S21.**

NNN and NN$_{PKK}$N$_{THK}$ did not form membrane-associated punctate dots without the expression of AVRblb2. (A) Hellfire-tagged Rpi-blb2 and NRC3s-GFP were co-expressed with copper-inducible flag-AVRblb2 and CUP2-P65. Water was infiltrated at 24 hpi. Samples were examined at 3, 6, 12, and 30 hours post-water treatment. Scale bars represent 10µm. SOBIR1-mcherry was used as a plasma membrane marker. (B) Quantification of punctate dots of NNN and NN$_{PKK}$N$_{THK}$ examined at 3, 6, 12, and 30 hours post-water treatment.

**Table S1.**
List of constructs used in cell death assays

| Vector backbone | Promoter | protein name | Tag | $OD_{600}$ | Reference |
|---|---|---|---|---|---|
| pICH86988 | 35S | (empty vector) | none | 0.5 | This study |
| pICH86988 | 35S | NRC3 variants | C terminal myc | 0.5 | This study |
| pk7wGF2 | 35S | Rpi-blb2 | N terminal GFP | 0.2 | (*1*) |
| pGWB12 | 35S | AVRblb2 | N terminal flag | 0.1 | (*1*, *2*) |
| pTFS40 | 35S | Pto | C terminal GFP | 0.5 | (*3*, *4*) |
| pT50 | 35S | AvrPto | C terminal flag | 0.2 | (*3*, *4*) |
| pBIN | | R1 | none | 0.2 | (*5*, *6*) |
| pk7wGF2 | 35S | Avr1 | N terminal GFP | 0.1 | (*5*, *7*) |
| pBIN | GPA2 | Gpa2 | none | 0.5 | (*8*) |
| pBIN | 35S | Rbp1 | none | 0.5 | (*8*) |
| pG3101 | 35S | Rx | C terminal HA | 0.05 | (*9*, *10*) |
| pG3101 | 35S | CP | C terminal CBP-SBP | 0.05 | (*10*) |
| pICH86977 | 35S | Sw5b | none | 0.2 | (*5*, *11*) |
| pICH86977 | 35S | Nsm | none | 0.1 | (*5*, *12*) |

**Table S2.**
List of constructs used in disease resistance assays

| Vector backbone | Promoter | protein name | Tag | $OD_{600}$ | Reference |
|---|---|---|---|---|---|
| pICH86988 | 35S | (empty vector) | none | 0.5 | This study |
| pK7WGR2 | 35S | Rpi-blb2 | N terminal RFP | 0.2 | (*1*) |
| pICH86988 | 35S | NbNRC4 | C terminal myc | 0.05 | (*5*) |
| pICH86988 | 35S | SlNRC3 | C terminal myc | 0.05 | This study |
| pICH86988 | 35S | NbNRC3 | C terminal myc | 0.05 | This study |
| pICH86988 | 35S | $NN_{PKK}N_{THKI}$ | C terminal myc | 0.05 | This study |

**Table S3.**
List of constructs used in co-IP assays

| Vector backbone | Promoter | protein name | Tag | $OD_{600}$ | Reference |
|---|---|---|---|---|---|
| pGWB12 | 35S | AVRblb2 | N terminal flag | 0.1 | (*1*, *2*) |
| pGWB555 | 35S | RFP | none | 0.2 | (*1*, *2*) |
| pK7WGR2 | 35S | Rpi-blb2 | N terminal RFP | 0.2 | (*1*) |
| pICH86988 | 35S | NbNRC3 | C terminal myc | 0.5 | This study |
| pICH86988 | 35S | $NN_{PKK}N_{THK}$ | C terminal myc | 0.5 | This study |

**Table S4.**
List of constructs used in TurboID-based proximity labeling assays

| Vector backbone | Promoter | protein name | Tag | $OD_{600}$ | Reference |
|---|---|---|---|---|---|
| pICH47742 | CBS4 | AVRblb2 | N terminal flag | 0.1 | This study |
| pICH47742 | CBS4 | AVRblb2 | N terminal GFP | 0.1 | This study |
| pGWB555 | 35S | RFP | none | 0.2 | (*1*, *2*) |
| pK7WGR2 | 35S | Rpi-blb2 | N terminal RFP | 0.2 | (*1*) |
| pICH47742 | 35S | CUP2-P65AD | | 0.2 | This study |
| pICH86988 | 35S | NbNRC3 | C terminal TurboID40-flag | 0.5 | This study |
| pICH86988 | 35S | $NN_{PKK}N_{THK}$ | C terminal TurboID40-flag | 0.5 | This study |

**Table S5.**
List of constructs used in cell biology assays

| Vector backbone | Promoter | protein name | Tag | $OD_{600}$ | Reference |
|---|---|---|---|---|---|
| pICH47742 | CBS4 | AVRblb2 | N terminal flag | 0.1 | This study |
| pGWB12 | 35S | AVRblb2 | N terminal flag | 0.1 | (*1*, *2*) |
| pT50 | 35S | AvrPto | C terminal flag | 0.2 | (*3*, *4*) |
| pGWB555 | 35S | RFP | none | 0.2 | (*1*, *2*) |
| pICH47751 | 35S | Rpi-blb2 | none | 0.2 | This study |
| pICH47742 | 35S | CUP2-P65AD | C terminal NLS | 0.2 | This study |
| pK7WG2 | 35S | NbSOBIR1 | C terminal mCherry | 0.2 | (*13*) |
| pICH86988 | 35S | $NbNRC4^{AAA}$ | C terminal GFP | 0.2 | This study |
| pTFS40 | 35S | Pto | C terminal GFP | 0.5 | (*3*, *4*) |
| pICH86988 | 35S | $NbNRC3^{L21E}$ | C terminal GFP | 0.5 | This study |
| pICH86988 | 35S | $SlNRC3^{L17E}$ | C terminal GFP | 0.5 | This study |
| pICH86988 | 35S | $N^{L21E}SS$ | C terminal GFP | 0.5 | This study |
| pICH86988 | 35S | $N^{L21E}N_3N_{T5be}$ | C terminal GFP | 0.5 | This study |
| pICH86988 | 35S | $N^{L21E}N_{PKK}N_{THK}$ | C terminal GFP | 0.5 | This study |

**Data S1. (Separate file)**
List of primers used in this study.

**Data S2. (Separate file)**
List of plasmids used in this study.

**Data S3. (Separate file)**
Sequences and accession numbers of NRC used in this study.

**Data S4. (Separate file)**
Raw data of the entropy analysis.

**Data S5. (Separate file)**
Raw data of FEL analysis and SLAC analysis.

**Data S6. (Separate file)**
Raw data of ancestral sequence reconstruction.
